# A multiscale modeling approach to study the role of mechanics and inflammation in pathophysiology of articular cartilage

**DOI:** 10.64898/2025.12.29.696945

**Authors:** Satanik Mukherjee, Raphaelle Lesage, Liesbet Geris

## Abstract

Mechanical loading regulates chondrocyte health in articular cartilage. While physiological stimuli maintain homeostasis, supra-physiological stimuli from joint injuries disrupt it, leading to osteoarthritis (OA). OA is a prevalent degenerative joint disease affecting millions worldwide. OA progression involves complex mechanical and biochemical interactions across multiple length scales, which are challenging to investigate experimentally. In silico models provide an effective framework to explore these mechanisms.

This study developed an integrated multiscale modeling framework for articular cartilage. It combined finite element (FE) models at tissue and cellular scales with an intracellular gene/protein regulatory network. The network incorporated key chondrocyte mechanotransduction and inflammatory pathways. A Hill’s function was used to link cellular forces from the FE model to a mechanical loading input to the regulatory network. Hill’s function constants were calibrated using a genetic algorithm approach. Calibration was performed by matching experimental and simulated expressions of COL-II and ADAMTS5 of cartilage explants under 20% dynamic compression.

As a validation step, model simulations were performed at 10% dynamic compression of cartilage explants. COL-II and ACAN were overestimated, and ADAMTS5 was underestimated compared with experimental data. Furthermore, predicted sGAG loss matched the trend of experimental data. Simulated chondrocyte responses for varying spatial locations revealed spatial heterogeneity of chondrocyte activity. Overall, the multiscale modeling workflow developed in this study provides a first step towards a powerful tool to increase the understanding of the complex interplay of mechanics and inflammation in articular cartilage. By integrating tissue, cellular, and intracellular scales, it offers a comprehensive framework for studying cartilage mechanobiology and guiding future therapeutic strategies.

**Highlights:**

- Developed an integrated multiscale model linking tissue, cellular mechanics, and gene regulation in articular cartilage.
- Coupled cellular mechanical forces to gene regulatory networks using Hill’s function approach.
- Calibrated Hill’s function parameters via genetic algorithm using experimental cartilage explant compression data.
- Predicted spatial heterogeneity of chondrocyte activity and cartilage biomarker expression under dynamic compression
- Established computational framework can be used for studying cartilage mechanobiology and osteoarthritis therapeutic strategies

## 1. Introduction

Mechanical loading is a critical factor that regulates the health of chondrocytes, the primary cells residing in articular cartilage. While physiological mechanical stimuli are essential for chondrocyte homeostasis, supra-physiological mechanical stimuli, such as those occurring during joint traumatic injuries, can disrupt stable cartilage homeostasis, leading to catabolic processes that contribute to osteoarthritis (OA) [1]. OA is the most prevalent degenerative joint disorder, affecting over 595 million people globally and contributing to significant disability and socioeconomic burden [2]. As populations age and obesity rates increase, OA prevalence is expected to reach 1 billion by 2050 [2].

Understanding chondrocyte mechanobiology provides insight into both cartilage degeneration and regeneration. Mechanical signal transduction in cartilage is inherently a multiscale process. Mechanical loads experienced by the joints during daily activities, propagate from the cartilage tissue to the chondrocytes. Chondrocytes sense mechanical stimuli through mechanosensitive receptors like integrins and ion channels such as TRPV4 and Piezo 1/2 [3–5], which initiate a cascade of intracellular biochemical processes via mechanotransduction pathways. Disruptions in these pathways can impair matrix maintenance, leading to early damage in the pericellular region that gradually extends into the extracellular matrix (ECM) [6].

In addition to being mechanosensitive, chondrocytes also respond to pro-inflammatory cytokines. Although OA is not traditionally considered inflammatory, low-grade synovitis and cytokine signaling (e.g., IL-1*β*, TNF-*α*) play critical roles in disease progression [7–10]. Moreover, mechanoregulatory and inflammatory signals are strongly interconnected in cartilage. Below a certain threshold, loading may dampen inflammation, but excessive loading can exacerbate it [11, 12].

Deciphering the crosstalk between mechanical and inflammatory cues is essential for understanding OA mechanisms and informing mechanotherapies or pharmacological treatments. However, studying this complex interplay experimentally is challenging due to the multiscale and multifactorial nature of cartilage regulation. *In silico* models provide a powerful platform to integrate these diverse mechanisms [13, 14]. In this context, previous *in silico* models of chondrocyte mechanotransduction were often linear, deterministic, or phenomenological regarding the intracellular component [11, 15], lacked detailed cell regulatory networks [16–18] and/or lacked a multiscale perspective [19–21].

The present study seeks to develop an integrated multiscale modeling workflow by combining a multiscale finite element (FE) model of articular cartilage with a mechanistic chondrocyte intracellular model. The intracellular model aims to capture relevant mechanisms for studying the interplay between mechano-regulatory and inflammatory pathways and their influence on chondrocyte phenotypic changes. In what follows, the different length scales involved in the multiscale model are described, along with explanations on connecting the length scales. A brief description of the biological mechanisms integrated into the intracellular regulatory networks is provided along with the main principles of its implementation. Subsequently, the calibration procedure using genetic algorithms based on *ex vivo* data reported in the literature is described. Finally, some validation activities for demonstrating model credibility of the multiscale model is presented.

## 2. Materials and Methods

### 2.1. General workflow of the multiscale modeling approach

The general multiscale modeling framework developed in this study consisted of three different *in silico* models, representing three length scales as shown in Figure 1. The models were mechanically coupled to each other using a post-processing approach [22, 23], such that the outputs from a larger scale were used as inputs to drive a smaller scale model. The general description of the models is provided below in descending order of length scales.

**Figure 1:**
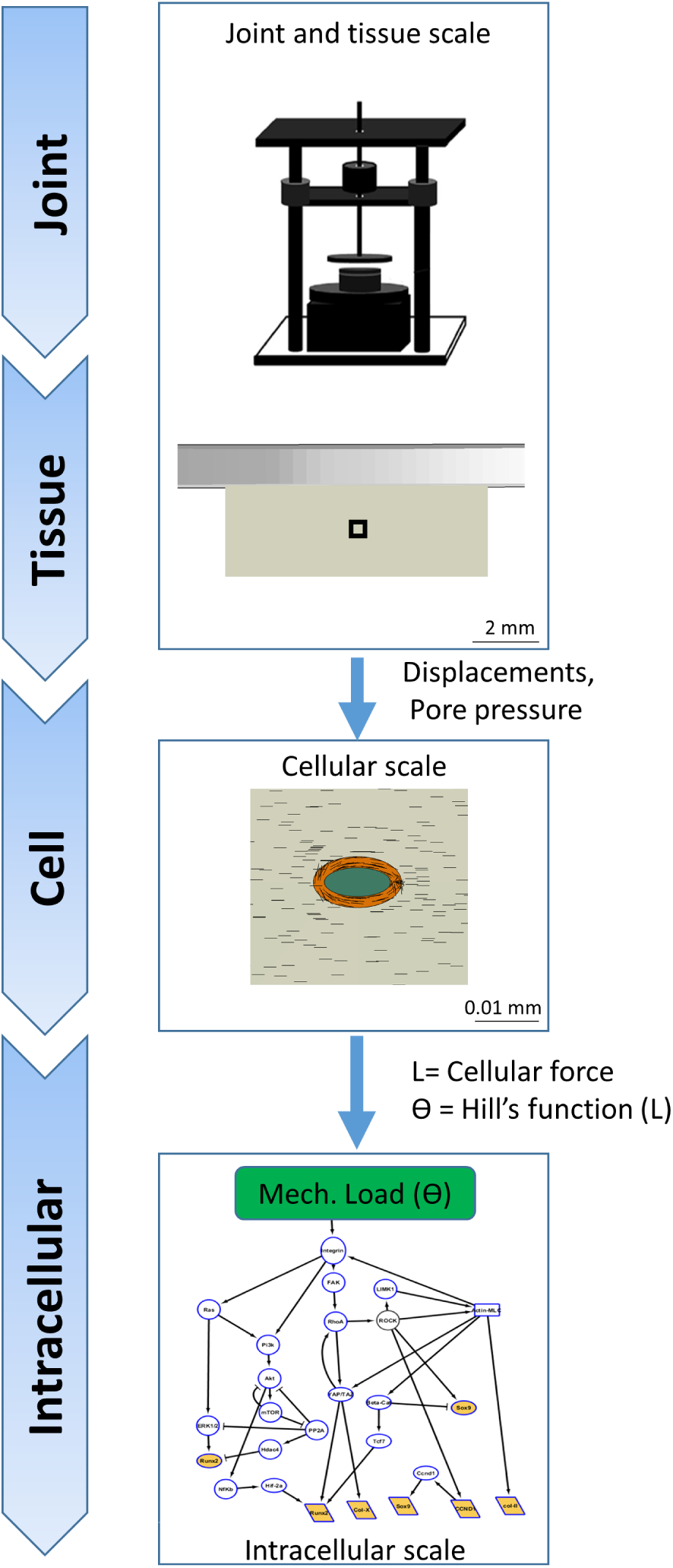
Overview of the multiscale model workflow showing the different length scales involved in an in vitro setup for cartilage compression. The black box in the tissue scale for the cylindrical cartilage explant indicates the location of the chondrocyte simulated in the cell level model. Black lines in the cellular scale denotes the fibril orientations in the ECM and PCM. The density of fibrils in the figure is reduced from the actual implementation for better visualization.*θ*=Mechanical loading input to networks, L=Cellular force from FE model

#### 2.1.1. Organ and tissue scale

The largest length scale of the multiscale model workflow comprises the organ and tissue scale. Organ is a general term used in the workflow that can encompass an *in vivo* case (such as the knee joint) or an *in vitro* setup that mimics the physiological mechanical loading in a joint. The tissue scale focuses on articular cartilage, which is the primary tissue of interest in this study.

Both the organ and tissue level consisted of an FE model that was developed for an *in vitro* setup as used in the study by Li et al. [12]. The setup consisted of an uniaxial dynamic compression bioreactor that imposed mechanical loading to the cartilage explant. The geometry and boundary conditions used in this FE model is described in detail in Section 2.2.

##### Constitutive model of articular cartilage

Articular cartilage was modeled as a fibril-reinforced poroviscoelastic (FRPVE) material, as described in previous studies [24, 25]. A detailed implementation of the constitutive model has been discussed in our previous study [26]. Briefly, this biphasic material model consists of a solid and a pore-fluid component, with the solid component further divided into fibrillar and non-fibrillar parts. The fibril structure was implemented as a network with two primary and 24 secondary fibril directions at each integration point. The non-fibrillar part of the cartilage, which primarily consists of proteoglycans, was modeled using a modified Neo-Hookean law, as described in [27]. The total stress at an integration point was given by:

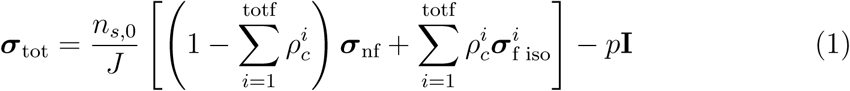

where **I** is the unit tensor, *n_s,_*_0_ is initial solid volume fraction, *J* is the determinant of the deformation tensor **F**, totf is the total number of fibrils considered at each integration point, *ρ^i^_c_* is the volume fraction of the *i^th^* collagen fibril with respect to the total volume of the solid matrix, ***σ***_nf_ is the stress in the non-fibrillar matrix, ***σ***_f iso_ is the stress in the collagen fiber network. The material model was implemented using the UMAT subroutine in Abaqus 2023 (SIMULIA, Providence, RI, USA)

Table 1 lists the values of the material parameters that were used for the simulations.

**Table 1:** Material parameters of the cartilage, extracellular matrix (ECM), pericellular matrix (PCM) and the chondrocyte used in this study(obtained from [28])

| Parameter | Cartilage/ECM | PCM | Chondrocyte |
| --- | --- | --- | --- |
| Shear modulus of the non-fibrillar matrix $G_m$ [MPa] | 0.9 | 0.9 | 0.017 |
| Shear modulus of the collagen matrix $G_{mf}$ [MPa] | 0.9 | 0.9 | - |
| Material constants: $E_1, E_2$ [MPa] | 4.316, 19.97 | 0.4316, 1.997 | - |
| Material constants: $K_1, K_2$ | 16.85, 41.49 | 16.85, 41.49 | - |
| Viscosity of collagen fibril $\eta$ [MPa-s] | $1.424 \times 10^5$ | $1.424 \times 10^5$ | - |
| Permeability constant $\alpha$ [$\text{m}^4/(\text{N-s})$] | $1.767 \times 10^{-15}$ | $1.767 \times 10^{-15}$ | $2.6 \times 10^{-10}$ |
| Nonlinearity term of permeability M | 1.339 | 1.339 | - |

#### 2.1.2. Connecting tissue scale to cellular scale

To mechanically couple the tissue scale model with the cellular scale model, a node-based sub-modeling technique was utilized in Abaqus. Given the biphasic material behavior of cartilage, the global displacements and pore pressure obtained from the tissue scale model simulation were applied as boundary conditions to drive the cellular sub-model. This post-processing method of connecting length scales enables simulations of multiple cellular scale models at different locations within the tissue scale model.

#### 2.1.3. Cellular scale

The cellular scale was modeled as a cube of edge length 100 *µ*m, consisting of a chondrocyte at its centroid, surrounded by its pericellular matrix (PCM) and extracellular matrix (ECM). To approximately satisfy mechanical consistency across spatial scales, the following considerations were made: firstly, ECM, which comprised 99.9% of the total volume of the cube, was assigned identical properties to those of the articular cartilage in the tissue-scale model. This choice was made to maintain consistency in the material properties between the ECM and the cartilage at the macroscale level. Secondly, the choice of a 100 *µ*m representative volume element (RVE) for the cube was based on previous studies in the literature [23, 29, 30]. This size was selected to ensure that the inclusion of the cell and PCM does not significantly alter the deformation field at the boundaries of the cube.

The 2.5 *µ*m thick PCM layer was modeled as an FRPVE material, with the 2 primary collagen fibrils oriented tangentially to the cell surface [31, 32] as shown in Figure 1. Since the diameter of type VI collagen fibrils, the predominant collagen type present in the PCM, is less than that of type II collagen fibrils predominantly found in the ECM [33], the collagen fibril stiffness in the PCM was assumed to be 10% of that in the ECM [22, 28].

The chondrocyte itself was modeled as a homogeneous, poro-hyperelastic solid using the same modified neo-Hookean material model as used to describe the mechanical behavior of non-fibrillar matrix of cartilage (section 2.1.1). The chondrocyte was modeled as an ellipsoid with its major axis parallel to the cartilage surface [34, 35]. The major and minor axis diameters were 15 *µ*m and 6.5 *µ*m respectively. The material parameters for the cell, ECM and PCM used in the study are shown in table 1. The ECM, PCM and chondrocyte were connected to each other using the ‘tie’ constraint in Abaqus, which ensured continuity of displacements and pore pressures across the ECM-PCM and PCM-cell interfaces. A mesh convergence study was performed for the cellular scale model as shown in Supplementary Figure S2

#### 2.1.4. Connecting cellular scale to intracellular scale

In the FE model at the cellular scale, the total force (L) transmitted to the cell via the PCM was determined by summing the magnitude of forces acting on the nodes of the cell that were connected to the PCM using the ‘tie’ constraint. However, the total force obtained could not directly be applied as the mechanical load input to the intracellular regulatory network in the multiscale modeling workflow. This was because the regulatory network is semi-quantitative in nature (with activity of each biomolecule representing a node in the network ranging from 0-1) whereas the cellular force calculated in the FE model was purely quantitative in nature (measured in newtons). To translate the cellular forces (L) calculated from the FE model to a mechanical loading input (Θ) to the intracellular network, a Hill’s function was chosen, as cellular mechanosensing is inherently nonlinear, saturable, and threshold-dependent[36]. This formulation, as shown below, provides a smooth, bounded mapping between cellular force magnitude (L) and mechanical loading input (Θ), converting physical forces into a non-dimensional value in the range [0,1] that is compatible with the regulatory network and can be calibrated using experimental data.

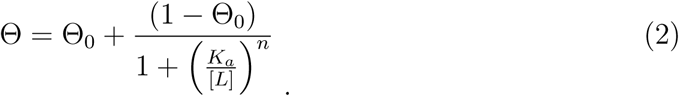

Θ_0_ represents the basal level of mechanical load sensed by the chondrocyte which is the mechanical load in the absence of any external mechanical stimulus. Since chondrocytes can detect ECM stiffness through focal adhesions via integrins [37], this basal load was assumed to be 0.1. L is the total force acting on the cell in the FE model, *K_a_* and n are constants for the Hill’s function. Physically, *K_a_* represents the magnitude of cellular force L, at which the mechanical loading input Θ is halfway between its basal value (Θ_0_ = 0.1) and its maximum possible value (which is 1). *K_a_*indicates the sensitivity of the cellular mechanosensors (primarily the integrins, as will be discussed in following section) to the mechanical force. A lower *K_a_* means the cell is more sensitive to smaller magnitudes of mechanical loading. The value of n affects how sharply the mechanical loading input function Θ changes as the L increases. The values of constants *K_a_* and n were calibrated using experimental data as discussed in Section 2.2

#### 2.1.5. Intracellular scale: gene and protein regulatory network

The intracellular pathways triggered by mechanical loading were modeled starting from a gene and protein regulatory network. In this network, nodes represent key biological components (proteins or genes) involved in chondrocyte biology. Directed edges between nodes signify interactions or activating/inhibitory influences between source and target proteins, or between transcription factors (TFs) and their target genes. The network is structured into two interactive layers: one focused on protein signaling and another on gene regulation. The protein signaling network layer spans from the binding of growth factors to their respective receptors, down to the TFs entering the nucleus. The gene regulatory network (GRN) layer illustrates how transcription factors regulate the expression of target genes, which in turn code for proteins involved in the signaling network. These two network layers are inter-connected, as each biological component is represented by both a gene in the GRN layer and its corresponding protein in the signaling network layer. Together, the two network layers form an interactive knowledge base for chondrocyte signaling.

The mathematical modeling of this network consisted of two distinct steps: first, constructing the network by representing the topological information of intracellular biochemical processes as a directed graph, and second, implementing a framework to simulate the dynamics of the network. An explanation of these two steps is provided below.

##### Construction of the gene and protein regulatory networks

The regulatory networks were constructed using both knowledge-driven (based on known biological principles) and data-driven approaches. The knowledge-based regulatory network was manually curated from the literature. It was complemented with data-driven network inference to uncover *de novo* regulatory links, reveal previously unstudied or undiscovered interactions, and minimize bias associated with human literature curation. A detailed methodology of the regulatory network construction can be found in Lesage et al. [38]. In the present study, extensions were made to the previously published regulatory network in Lesage et al. [38], to include additional biochemical mechanisms related to mechanotransduction, inflammatory pathways, and their crosstalk in chondrocytes.

###### Mechanotransduction pathways

In the modeled pathways, mechanical load (*θ*) is transduced into biochemical cascades by integrins and focal adhesion kinase (FAK) complexes, activating RhoA/ROCK signaling that regulates actin cytoskeleton assembly and contractility, which influences cell differentiation and response to mechanical stimuli. This pathway also affects transcription factors like SOX9, which promotes ECM production, and YAP/TAZ, which regulate chondrogenic phenotypes and prevent hypertrophy. Additionally, the mTOR pathway is incorporated, linking integrins to PI3K/AKT/mTOR signaling, which regulates factors like GLI2 and RUNX2, contributing to chondrocyte mechanosensitivity.

In addition to integrins, chondrocytes also rely on mechanosensitive ion channels for mechanosensing and mechanotransduction, collectively referred to as the ‘channelome’ [39]. This network included the cation channel TRPV4 (transient receptor potential vanilloid 4), that plays a key role in the chondrocyte’s metabolic response to dynamic mechanical loading [4]. Recent research [40] suggests that TRPV4 activation in chondrocytes occurs due to osmotic fluctuations caused by the deformation of the chondrocyte’s osmotically active microenvironment. Since the exact mechanism linking deformation, osmotic changes and TRPV4 activation is very complex and not fully established in literature, it was assumed that the mechanical load (*θ*) on the chondrocytes was directly responsible for TRPV4 activation. This assumption was based on the fact that mechanical load is directly proportional to the deformation of the microenvironment, as chondrocytes are directly attached to the extracellular matrix via focal adhesions. Supplementary data S1.1 contains a detailed explanation of the mechanotransduction pathways that were included in the current study.

###### Inflammatory pathways

Pro-inflammatory cytokine signaling (e.g., IL-1*β*, TNF-*α*) is known to play a critical role in OA progression [7–10]. In this network, we simplified pro-inflammatory signaling by representing key mediators (e.g., IL-1*β* or TNF-*α*) with a single *‘pro-inflammatory cytokines’* node that activated an inflammation receptor (‘R-inflam’). This was an important simplification allowing us to focus on the intracellular interplay between mechanotransduction and inflammation rather than the role of specific receptors and signaling molecules. In the network, the *‘pro-inflammatory cytokines’* node induces matrix degradation, suppresses matrix biosynthesis, and amplifies its own production through feedback loops involving signaling cascades like TAK1, NFKB, and MAPKs. These pathways include activation of iNOS, which further fuels inflammation, and Wnt signaling, which also contributes to NFKB activation. Negative feedback loops, including SOCS proteins and the inhibition of TAK1, help regulate this auto-inflammatory response, with IGF signaling playing a role in modulating inflammation as well as promoting tissue repair. Supplementary data S1.2 contains a detailed explanation of the inflammatory pathways that were included in this study.

###### Mechano-inflammatory crosstalk pathways

Pathways involved in mechanotransduction and inflammation are interconnected, as shown by Madhavan et al. [41], who found that cyclic tensile strain suppresses pro-inflammatory gene induction by IL-1B, TNF-A, and LPS. In the current model, we included several direct molecular mechanisms linking mechanical loading and inflammation. For instance, integrin downstream signaling, particularly through *α*_5_*β*_1_, upregulates MMP13 expression in chondrocytes when combined with inflammatory cues like IL-1B. Additionally, the model accounts for the interaction between YAP/TAZ and NFKB, where mechanical signals at physiological levels inhibit NFKB activation. Finally, ECM degradation products (DAMPs) activate inflammatory cytokine receptors and integrins, while mechanosensitive calcium signaling (TRPV4/Ca2+) inhibits ADAMTS5, highlighting the complex crosstalk between mechanical and inflammatory pathways. Supplementary data S1.3 contains a detailed explanation of the mechano-inflammatory crosstalk pathways that were included in this study.

The complete network consists of 74 nodes and 299 interactions. Important interactions and crosstalks related to the mechanotransduction and inflammatory pathways are reported in an activity flow diagram for illustration purposes in Figure 2, providing an incomplete view of the full interactome integrated in the model.

**Figure 2:**
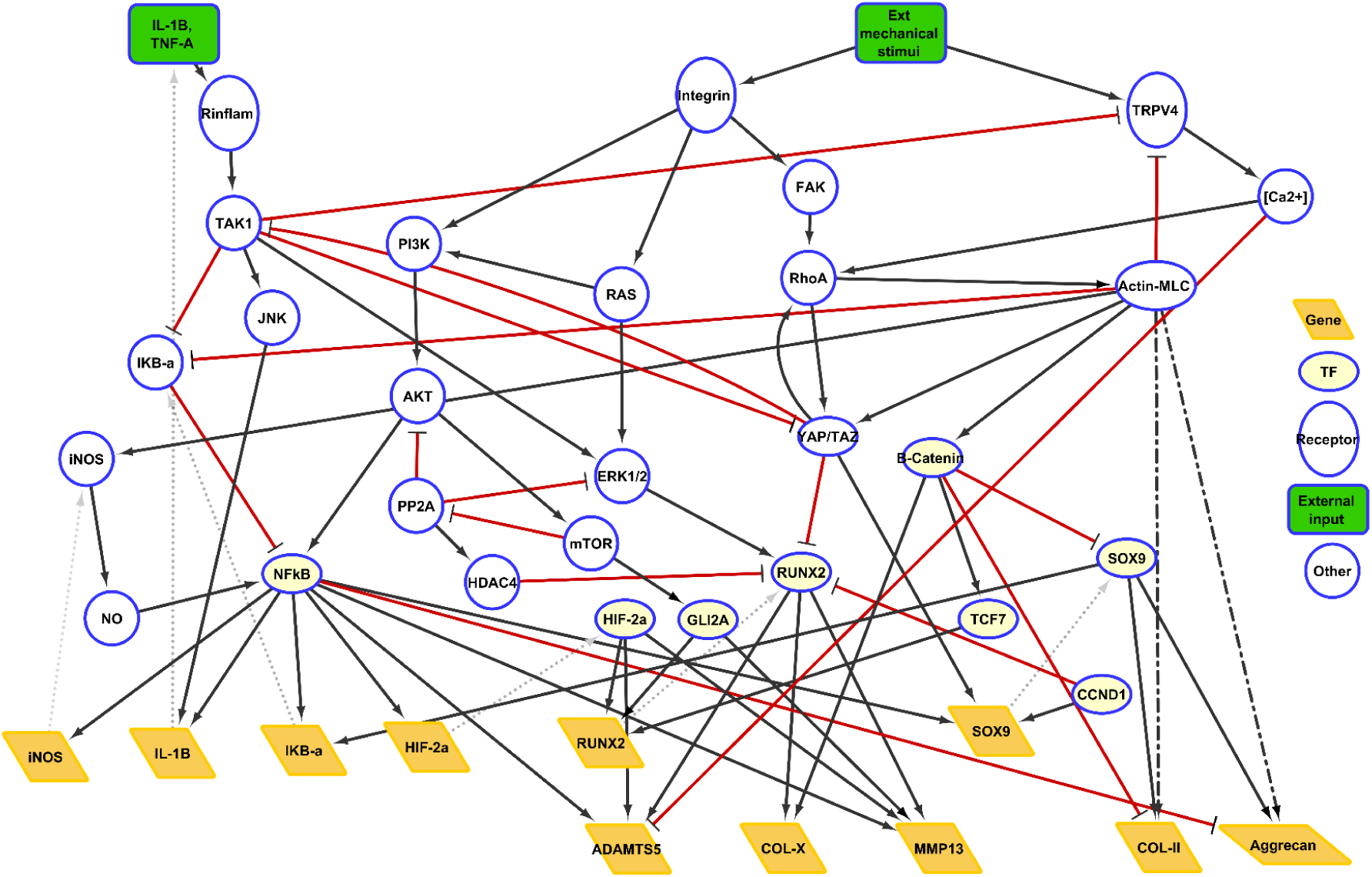
Snapshot of the mechanotransduction and pro-inflammatory signaling and their crosstalk in the in silico chondrocyte. This activity flow diagram is a selection of important crosstalks from the model and related to mechanotransduction or inflammation, for illustration. External cues (in green) are either the mechanical loading or pro-inflammatory cytokines. The signal is transduced through signaling molecules (‘Other’) up to transcription factors (‘TF’ in blue circled yellow ellipse), which up- or down-regulate the expression of downstream genes (yellow parallelograms). As a result, genes increase the amount of the corresponding upstream proteins (grey dotted lines). Black arrows (resp. red T-ended lines) represent activation (resp. inhibition) interactions or influences. Influences may be direct or indirect in chondrocytes, the dashed black lines represent interactions that involve an intermediary node in the model but that is unrepresented in the graph.

##### Simulating the dynamics of the regulatory networks

The developed regulatory network was translated into a mathematical model using the additive formalism with priority classes developed by Kerkhofs et al. [42]. In this approach, biological interactions were considered to occur on two distinct time scales: the fast (protein signaling) and the slow (gene regulation) levels. Consequently, each biological entity or node in the network could have both fast and slow upstream regulators.

The **global activity** of a node is defined as the product of its gene expression level and the activation level of the corresponding protein. Biologically, this measure reflects the node’s overall regulatory impact on downstream neighbors, capturing both transcriptional activity and functional protein activity. The activity of a node at the gene and protein levels is determined by summing the effects of upstream activators, which enhance the node’s activity, and subtracting the effects of inhibitors, which reduce it.

A Monte Carlo canalization analysis was performed to identify stable states of the system under mechanical loading. The mechanical loading input (*θ*) was constrained, while other nodes were updated freely across 10,000 random initializations. Among multiple attractor states identified, the one exhibiting healthy chondrocyte behavior (>0.7 activity for SOX9, COL-II, and ACAN) was selected as the baseline healthy state for subsequent perturbation analyses.

In order to simulate different mechanical loading and/or cytokine conditions, perturbations of the corresponding node from the healthy state were applied for an indefinite period of (computational) time steps and the new phenotypes were assessed after the system had settled in stable states under the perturbed environment. During the perturbation, the system could settle in slightly different final states because of the stochastic nature of the model. To account for this variability, each perturbation simulation was repeated 100 times. A weighted average of the resulting final profiles was then computed, with weights proportional to the frequency of each outcome.

#### 2.1.6. Implementation of the multiscale model

The development and analysis of FE models were performed using Abaqus 2023. The primary collagen fibril orientations in the ECM/PCM of the cell, were determined using scripts developed in-house in Matlab R2022b (The MathWorks Inc., Natick, Massachusetts). Subsequently, to implement the fibril reinforced poroviscoelastic model of cartilage, ECM and PCM, multiple user-defined subroutines in Fortran were used. In order to run simulations in parallel across multiple cores, existing UMAT subroutines were made thread-safe by removing common blocks, and UEXTERNALDB subroutines were developed to implement collagen fibril orientations. These improvements allowed FE simulations to be run in parallel, significantly reducing computation time. All FE simulations were performed on the wICE high-performance computing (HPC) cluster at the Flemish Supercomputing Center (VSC), KU Leuven. The cluster consists of 172 IceLake nodes, each equipped with two Intel Xeon Platinum 8360Y CPUs (2.4 GHz, 36 cores per CPU, 1 NUMA domain, and 1 L3 cache per CPU), 256 GiB RAM, and 960 GB SSD local disk. The default memory per core is 3400 MiB. Job submissions were managed via Slurm scripts. For the simulation of intracellular gene/protein regulatory network, in-house Matlab scripts were developed.

### 2.2. Calibration and validation of the multiscale model with in vitro experimental data

Important steps in establishing the credibility of multiscale models are their calibration and validation with experimental data. As described in Section 2.1, the FE models at individual length scales already used validated material parameters from the literature, and the regulatory network was developed from experimental data and its predictive value investigated by comparison with *de novo* experiments. However, the constants for the Hill’s function, used to translate cellular forces (L) on the cellular-scale FE model into mechanical loading input (*θ*) for the regulatory network, still required calibration with experimental data. Furthermore, the overall predictive capacity of the multiscale model also required validation against independent experimental observations.

For this calibration, an *in vitro* dynamic compression study of cartilage explants by Li et al. [12] was chosen, which applied varying levels of dynamic compression (10% strain to 30% strain) to bovine cartilage explants. This study also investigated the combined effect of dynamic compression and pro-inflammatory cytokines(combination of TNF-*α* (25 ng/ml), IL-6 (50 ng/ml) and sIL-6R (250 ng/ml)) on expression of relevant genes (such as ADAMTS5, COL-II, ACAN), making it suitable for calibrating our multiscale model. Additionally, the experiments measured the loss of sulfated glycosaminoglycan (sGAG), which provided an independent dataset to validate the overall multiscale model predictions.

#### 2.2.1. Multiscale digital twin of the in vitro bioreactor setup

A multiscale digital twin of the experimental setup was developed, that mimics the mechanical and biochemical environment experienced by the explants in the bioreactor (Figure 1). The digital twin incorporated three distinct length scales, with specific details for each scale outlined below.

i) **Organ and tissue scale:** an FE model of the cylindrical cartilage explant of 3 mm diameter and 1 mm thickness was created in Abaqus. The model was meshed with C3D8P elements and a mesh convergence study was performed to determine the optimum mesh density. FRPVE material properties as described in section 2.1.1 were assigned to the cartilage explant, with the primary collagen fibers parallel to the flat surface since the explants were harvested with the superficial zone intact. To simulate unconfined dynamic compression, the vertical displacement of the bottom surface of the explant was fixed using a YSYMM boundary condition. A zero pore pressure boundary condition was applied to the side of the cylinder to simulate free flow of pore fluid across this surface. Furthermore, displacement of two diametrically opposite points on the x and z axes at the bottom surface was constrained in the z and x direction, respectively, to prevent any rigid body motion along the x-z plane without compromising on the mechanics. Dynamic compression was applied to the explant using a rigid disc with a diameter of 8 mm, modeled as a discrete rigid body in Abaqus and meshed with R3D4 elements. The loading protocol for dynamic compression was based on the experiment by Li et al. [12], and was applied to a reference point on the rigid disc. First, a pre-strain of 10% of the cartilage thickness was applied over 1 second, and the strain was held constant for an additional 10 seconds to allow for stress relaxation. Following this, dynamic compression of 10% or 20% was applied using a haversine waveform (0.5 Hz, 40% duty cycle).
ii) **Cellular scale:** the cellular scale model as described in section 2.1.3 was positioned at the center of the cartilage plug, 0.5 mm from its top surface. The central position of the cell in the plug was chosen to capture an intermediate mechanical behavior that would be representative of the whole plug, given the gradient of mechanical stimuli present across its thickness.
iii) **Intracellular module:** the intracellular gene and protein regulatory network was applied as described in section 2.1.5.

#### 2.2.2. Sensitivity analysis of the regulatory network

In the experimental setup described in Section 2.2, cartilage explants were subjected to dynamic compression in the presence of pro-inflammatory cytokines. To ensure consistency between the experimental cytokine stimulus and its *in silico* representation in the multiscale model, a sensitivity analysis of the regulatory network was performed. This analysis determined the magnitude of the non-dimensional pro-inflammatory input that produced catabolic responses comparable to those observed experimentally. Sensitivity study of the regulatory network was performed using a two-factor full factorial design, by perturbing the pro-inflammatory cytokine and mechanical loading (*θ*) nodes of the network from 0 to 1 in steps of 0.1 (total of 121 simulations).

#### 2.2.3. Optimization using Genetic Algorithm approach

While the sensitivity analysis enabled mapping of the pro-inflammatory cytokine stimulus into a corresponding non-dimensional inflammatory input for the regulatory network, the mechanical loading input (*θ*) still required calibration. Specifically, the parameters of the Hill function *K_a_* and n (as described in section 2.1.4) used to translate cellular force L to the mechanical load input *θ* needed to be estimated to ensure accurate coupling between the FE-derived cellular forces (L) and the intracellular mechanotransduction pathways. These parameters were calibrated by minimizing the difference in gene expression levels of COL-II and ADAMTS5 between the experimental data [12] and the multiscale model under 20% dynamic compression and pro-inflammatory cytokine stimulation of the cartilage plug. Based on sensitivity analysis performed in section 2.2.2, a value of 0.695 was used to perturb the cytokine node of the network. The peak cellular force (L) from the 3rd cycle of dynamic compression was used to calculate the mechanical loading input (*θ*) to perturb the intracellular pathways. This value represented an intermediate cellular force, between the higher force of the first cycle and the lower steady-state forces of subsequent cycles. The minimization was performed using a genetic algorithm approach in Matlab. In the experiments by Li et al. [12], gene expressions were normalized with respect to a control case with no external loading and no cytokine stimulus. For consistency, the *in silico* gene expressions were also normalized to the control condition, which assumed a zero external load condition (L=0; *θ*=0.1) and no cytokine input to the regulatory networks. The objective function (f) for the optimization was as follows:

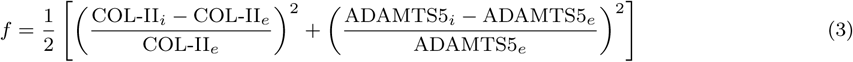

Subscripts *i* and *e* are for *in silico* and experimental values respectively. To prevent bias in the optimization due to differing gene expression magnitudes, each gene expression difference in the objective function was normalized by its corresponding experimental value, ensuring equal contribution from each gene.

#### 2.2.4. Validation of the multiscale model

Validation of the overall predictions of the calibrated multiscale model was performed in three steps by comparing model predictions with the corresponding experimental measurements in Li et al. [12]. First, ACAN gene expression was compared for the 20% strain condition. Next, the gene expressions of ACAN, COL-II, and ADAMTS5 were compared for the 10% strain condition. In both cases, cytokine treatment was applied. Finally, cumulative sGAG loss from cartilage explants to the medium after 8 days of treatment was compared with the numerically calculated aggrecan loss under the corresponding culture conditions. sGAG loss was compared for the unloaded, 10% and 20% strain conditions, under cytokine treatment. The numerical value of aggrecan loss for each condition was calculated as difference in global activity of the proteoglycan variable between the culture condition and the corresponding value at the baseline stable chondrocyte attractor.

### 2.3. Simulation of chondrocytes at varying spatial locations

Using the calibrated multiscale model, the biological response of cells at six distinct locations within the explant was assessed for 10% and 20% strain (Figure 6(a)). This involved simulating the cellular scale model at each location driven by the results from the tissue scale model, followed by the intracellular scale model. This analysis was performed to evaluate how spatial variations in mechanics in the explant influence cellular biological responses.

## 3. Results

### 3.1. Multiscale mechanical behavior of cartilage explants revealed by FE analysis

FE analysis of the cartilage explants revealed elevated fluid flow velocities along the curved (radial) surface of the explants (Figure 3(a,b)). The distribution of Von Mises stress exhibited a radial gradient, with highest stresses near the center of the explant under both 10% and 20% loading conditions (Figure 3(c,d)).

**Figure 3:**
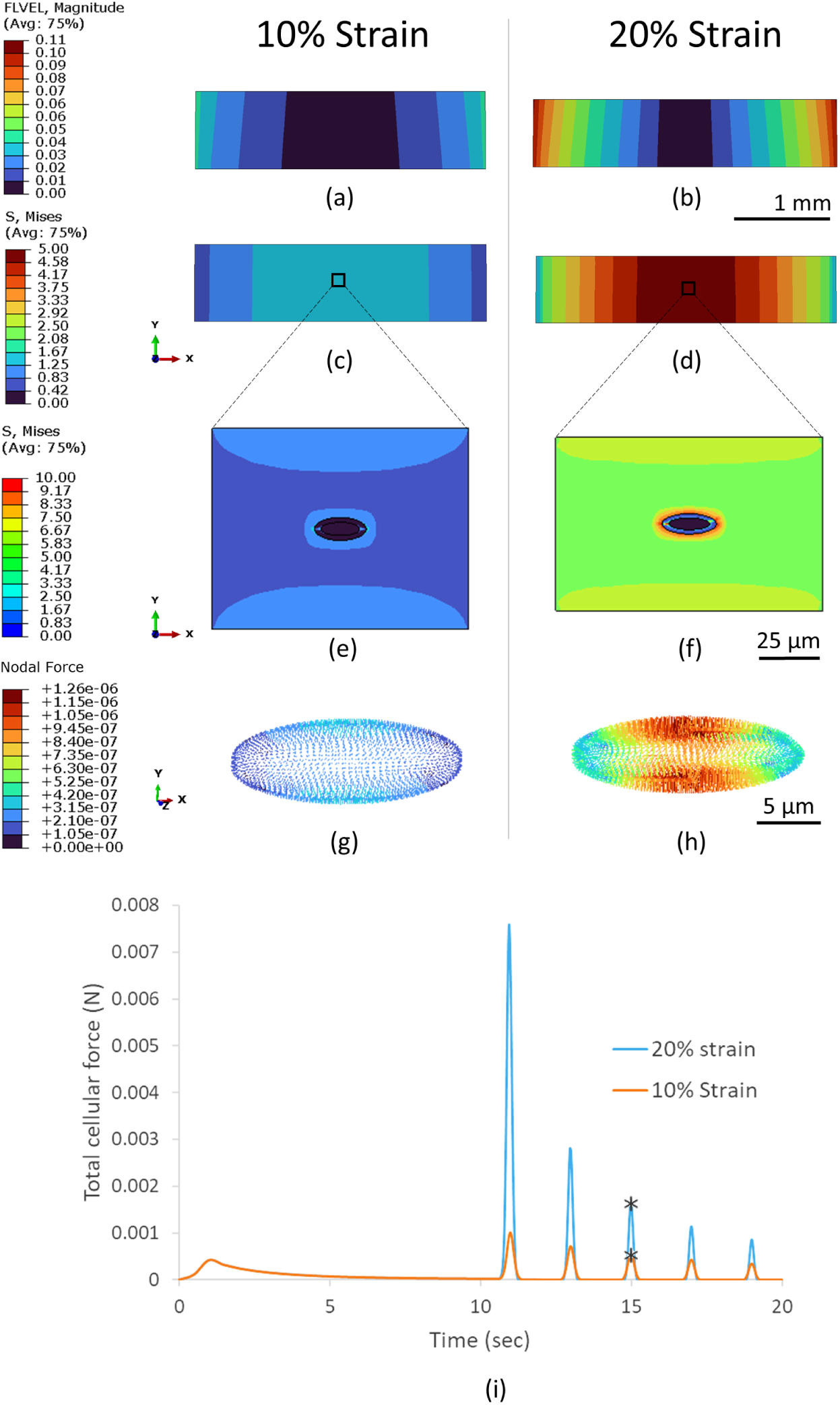
Results of FE analysis of the tissue and cellular scale mechanics. (a,b) Fluid velocity in cartilage explant. (c,d) Von Mises stress in cartilage explant. (e,f) Von Mises stress in cellular-scale model. (g,h) Nodal forces on the chondrocyte. (i) Total force variation on the chondrocyte. Black stars in (i) indicate the peak force during the third compression cycle, which was used as input for the intracellular model. All results shown for 10% and 20% strain respectively at the third cycle of compression.

At the cellular scale, FE analysis identified localized stress concentrations at the interfaces between the ECM-PCM and the PCM-cell (Figure 3(e,f)). The highest Von Mises stresses were found in the ECM adjacent to the PCM, followed by the PCM and then the cell itself. This trend corresponds to the relative stiffness of these components, with the ECM being stiffest and the cell the most compliant.

Nodal force analysis on the cells (Figure 3(g,h)) showed that the maximum forces were concentrated at the top and bottom regions of the cells. Additionally, the total force experienced by the cells demonstrated a decreasing trend from the first to the fifth compression cycle (Figure 3(i)), which can be attributed to the poroviscoelastic stress relaxation at tissue- and cell-levels.

### 3.2. Sensitivity analysis of the network reveals a complex interplay of mechanical and pro-inflammatory mediators

The sensitivity analysis of the network revealed a biphasic relationship between mechanical loading and the global activity of key cartilage matrix proteins, COL-II and ACAN, for pro-inflammatory cytokine levels below 0.7 (Figure 4). Specifically, mechanical loading (*θ*) in the range of 0.3 to 0.8 resulted in high global activity of ACAN and COL-II. However, when mechanical loading was either below 0.3 or above 0.8, the activity of these matrix proteins decreased.

**Figure 4:**
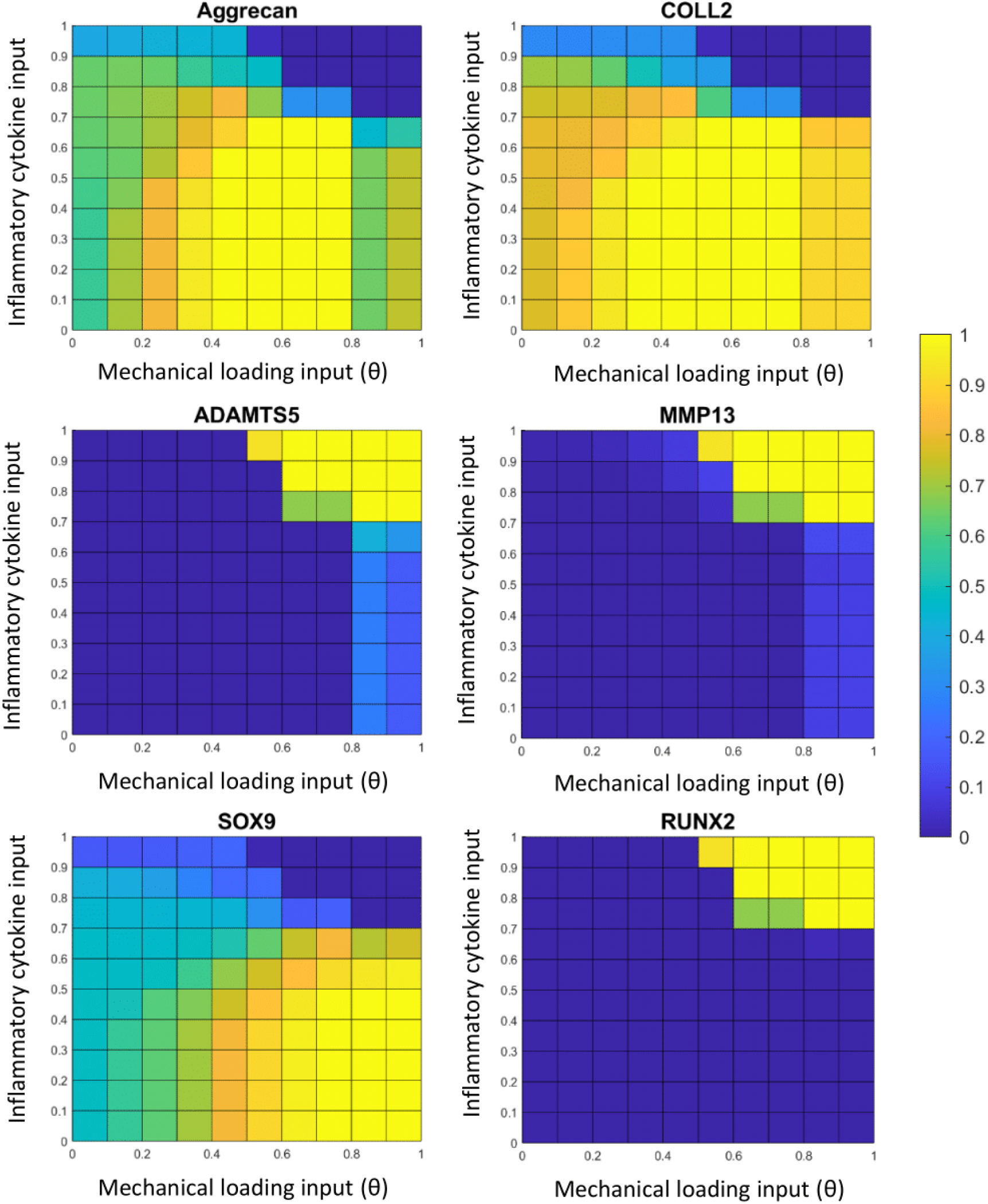
Results of the two-way sensitivity analysis of the intracellular network model. The x-axis corresponds to variations in the mechanical loading input (*θ*), and the y-axis corresponds to variations in the pro-inflammatory cytokine input. Each heatmap illustrates the resulting global activity of relevant biomarkers (in the range [0,1]) under the combined effects of mechanical loading and cytokine stimulation.

At higher cytokine levels (greater than 0.7), mechanical loads above 0.5 significantly reduced the global activity of ACAN and COL-II, while catabolic enzymes ADAMTS5 and MMP13 increased significantly, demonstrating a synergistic effect of high load and inflammation.

For cytokine levels below 0.7, the global activity of SOX9 increased monotonically with the increase in mechanical loading. However, when cytokine levels exceeded 0.7, mechanical loads greater than 0.5 led to a marked reduction in SOX9 activity. The global activity of RUNX2, a marker for pro-hypertrophic chondrocytes, was negligible under most conditions, except when both cytokine levels and mechanical loading were high.

### 3.3. Calibrated in silico multiscale model captures the gene expressions of COL-II and ADAMTS5

The calibrated values of Ka and n for the Hill’s function were 0.001656 and 1.870793, respectively, obtained at the 72nd generation of the genetic algorithm. Comparison of optimized numerical gene expression with experimental data for the 20% strain condition showed good agreement for ADAMTS5 and COL-II, with numerical predictions corresponding to approximately 0.7-fold and 1.2-fold of the experimental values, respectively (Figure 5). In contrast, ACAN expression was higher in the numerical model for the 20% strain condition, with numerical prediction corresponding to approximately 2.6-fold of the experimental values.

**Figure 5:**
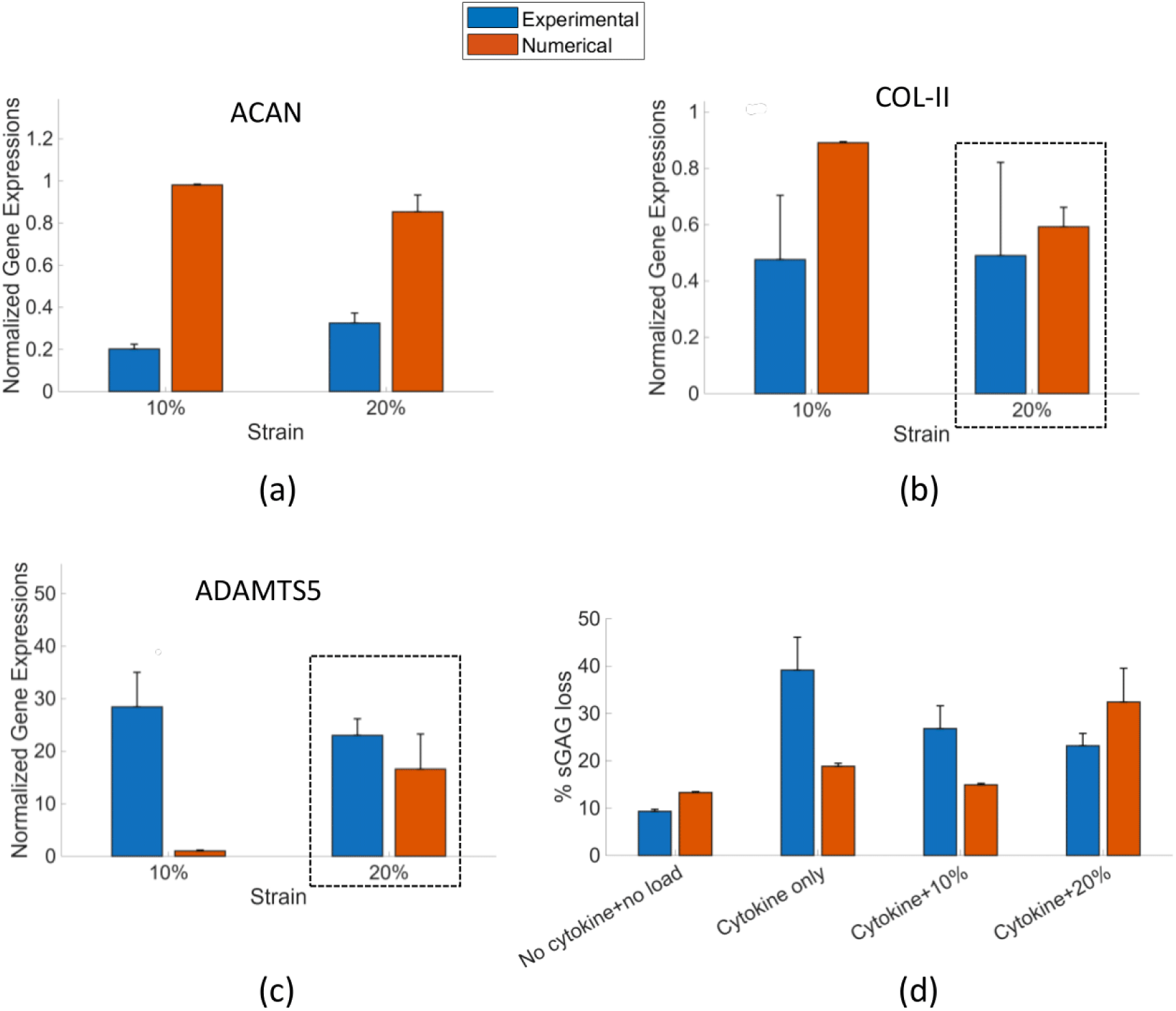
Results of the calibration and validation of the multiscale model. (a-c) Normalized gene expressions of ACAN, COL-II, ADAMTS5 under 10% and 20% dynamic compression of cartilage explants in the presence of cytokine stimulation. Gene expressions were normalized with respect to the no cytokine, no compression control condition which had an expression level=1. The dotted lines indicate the conditions included in the calibration, and the rest of the conditions were used for validation, (d) additional validation of the multiscale model by comparing the sGAG loss measured in experimental studies with the predicted value of Aggrecan loss from the model. All data are represented as mean *±* 95% confidence interval.

For the 10% strain condition, which served as an independent dataset for model validation, the numerical model predicted elevated COL-II and ACAN expression, equal to 1.9-fold and 4.9-fold of the experimental values, respectively. Conversely, ADAMTS5 expression was underpredicted at 10% strain, with the numerical prediction corresponding to approximately 0.04-fold of the experimental value.

### 3.4. In silico multiscale model predicts sGAG loss from cartilage

To further assess the predictive capability of the numerical model, sGAG loss from cartilage explants under different mechanical loading conditions with cytokine treatment was analyzed. The results, shown in Figure 5(d), indicated that the model predicted a 3.99% higher sGAG loss for the “no load, no cytokine” condition and a 9.25% higher sGAG loss for the “20% dynamic loading with cytokines” condition, than that observed experimentally. In contrast, sGAG loss was predicted to be 20.32% lower in the model for the “no load with cytokine” and 11.90% lower for the “10% dynamic loading with cytokine” condition as compared with experimental data. Both the numerical model and experimental data showed a consistent trend from “no load, no cytokine” to “no load with cytokine” and then to “10% dynamic loading with cytokines”. Specifically, sGAG loss increased upon cytokine treatment at no load and subsequently decreased under 10% dynamic loading. However, at 20% dynamic loading with cytokines, the numerical model predicted an increase in sGAG loss compared to the 10% strain case. Contrarily, the experimental data showed a continued reduction of sGAG loss, with the lowest sGAG loss observed at this condition.

### 3.5. Multiscale model captures location-specific cellular mechanobiological response

For 10% strain condition, global activity of biomarkers showed negligible differences between cells at the 6 different locations (Figure 6). However, for 20% strain, higher values for activity of anabolic markers like ACAN, COL-II and SOX9 were observed for the cells that were off-center (cells 2, 4 and 6) as compared to cells along the central axis (cells 1, 3 and 5). Correspondingly, the catabolic markers ADAMTS5, MMP13 and RUNX2 followed an opposite trend with higher values observed for cells along the central axis. Depth dependent effects were observed for 20% strain condition, with the catabolic markers ADAMTS5, MMP13 and RUNX2 showing slightly lower activity in cell 1, which was located centrally and closer to the loaded surface.

**Figure 6:**
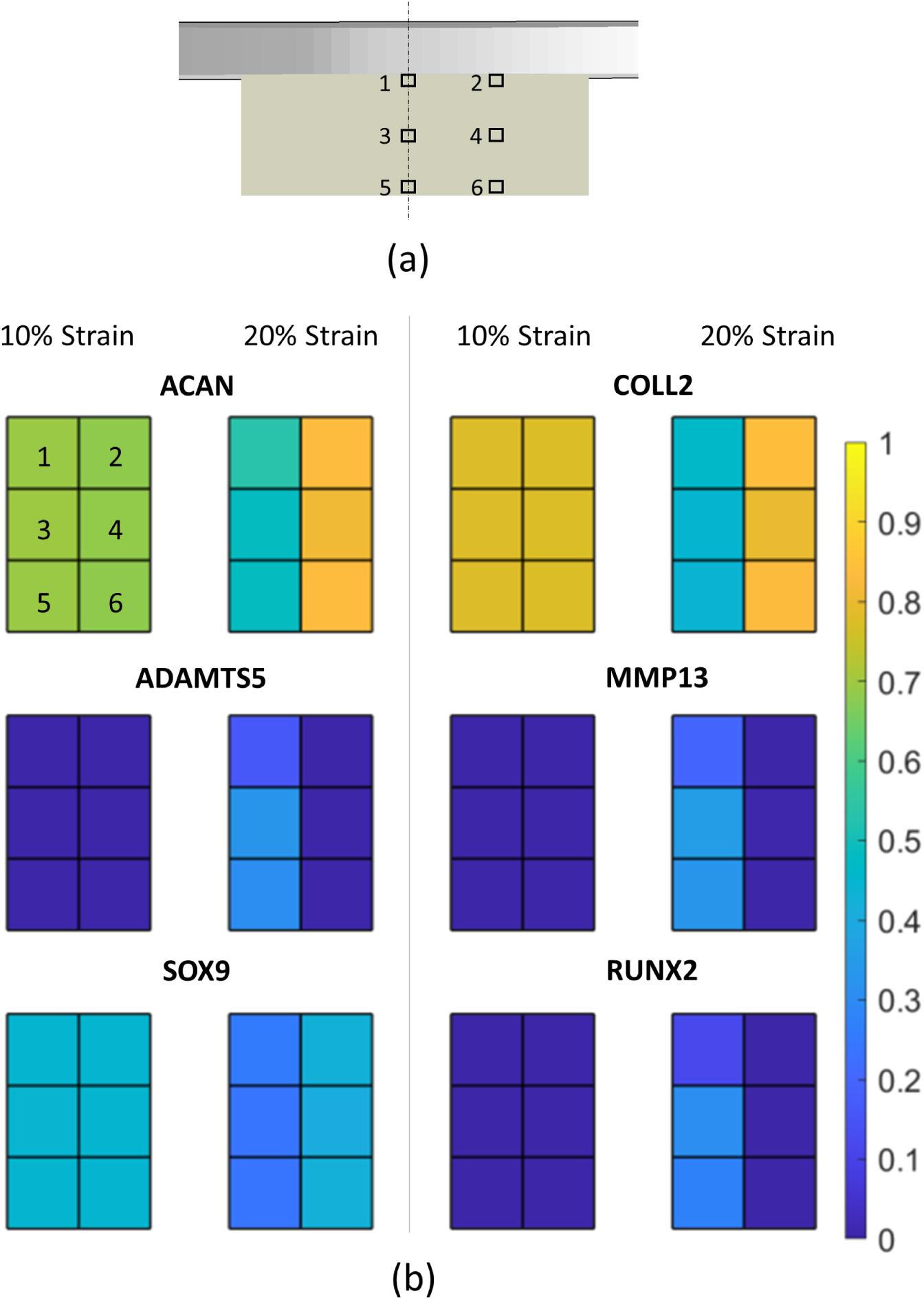
Simulation of 6 distinct spatial cell locations in the cartilage explant using the multiscale model. (a) Schematic representation of the cartilage explant showing six simulated spatial locations of individual cells, indicated by black boxes and labeled with corresponding cell numbers (1–6). (b) Heatmaps showing the global activity of ACAN, COL-II, ADAMTS5, MMP13, SOX9 and RUNX2 under 10% and 20% mechanical strain. Each heatmap is arranged in a 3×2 grid, with individual boxes numbered 1–6 corresponding to the cell locations defined in panel (a). Each box represents the global activity of a biomarker in the corresponding cell.

## 4. Discussion

In this study, a multiscale modeling workflow was developed for articular cartilage by coupling the mechanics at multiple length scales (from tissue to cell) with an intracellular gene/protein regulatory network. While there are studies in the literature that individually study either the multiscale mechanics of cartilage [22, 23, 29] or intracellular regulatory networks for cartilage [19–21, 38, 42], the present work uniquely integrates these two modeling paradigms into a single framework. This integrated framework enables direct investigation of how mechanical loading at the tissue and cellular scales propagates to intracellular regulatory responses, providing mechanistic insight into cartilage mechanobiology that cannot be obtained from either approach alone. In the tissue and cellular scale finite element models, a fibril-reinforced poroviscoelastic constitutive model [24] was used to describe the mechanical behavior of bulk cartilage, as well as the PCM and ECM. The simulations were run in parallel using multiple cores on an HPC cluster to reduce computational time. The modeling workflow was further calibrated and validated with respect to independent experimental data from the literature.

Sensitivity analysis of the intracellular regulatory network revealed how inflammation and mechanical loading interact to regulate relevant biomarkers. At low inflammatory levels (<0.7), the matrix proteins COL-II and ACAN exhibited a biphasic response to mechanical loading, while matrix-degrading enzymes ADAMTS5 and MMP13 increased only under high mechanical loads (>0.8), highlighting their role in reducing matrix proteins under excessive stimulation. At low mechanical loads (<0.3), reduced ACAN and COL-II activity reflected diminished matrix synthesis rather than increased degradation, consistent with lower activation of the anabolic transcription factor SOX9. This aligns with experimental observations of reduced sGAG synthesis at low mechanical loading in bovine cartilage explants by Li et al. [12]. At higher inflammation levels (>0.7), catabolic markers were markedly upregulated at moderate to high loads (>0.5), explaining the pronounced reduction in matrix proteins.

It should be noted that the values of biomarkers in the regulatory network are non-dimensional (range [0,1]). However, the Hill’s function implemented in the multiscale workflow provide a mechanistic link, allowing quantifiable cellular forces to be mapped to non-dimensional values for mechanical loading (*θ*) in the network. Overall, the sensitivity analysis demonstrated that the intracellular regulatory network within the multiscale framework can mechanistically capture the interplay between anabolic and catabolic pathways under varying inflammatory and mechanical conditions, providing a valuable tool to interpret how cellular forces might influence chondrocyte homeostasis.

This study included integrin-mediated mechanosensing as a primary mechanism of mechanical signal transduction in chondrocytes, as it is a well-studied pathway with substantial evidence supporting its role in mechanosensing [3, 43]. Although integrins comprise multiple subunits, limited evidence distinguishing subunit-specific downstream signaling motivated the assumption of a single representative integrin node in the regulatory network. In reality, integrin-mediated focal adhesions are localized to specific regions of the cell and exhibit complex binding dynamics [44]. However, we assumed a uniform attachment of the cell to the PCM using a ‘tie’ constraint in Abaqus. This assumption is supported by previous studies [45], in which immunofluorescent staining demonstrated a relatively uniform distribution of integrins around the chondrocyte.

Recent studies in the literature have shown the importance of Piezo channels (especially Piezo 1 and 2) as another mechanosensor for the chondrocyte, especially for high-strain scenarios [5, 46–48], which are not incorporated in the present study. In the future, as more downstream pathways of Piezo channels are delineated, incorporating Piezo channel-mediated mechanotransduction pathways could be a valuable addition to the present multiscale workflow. Furthermore, future studies could also incorporate the transport of nutrients, growth factors, and inflammatory cytokines across the cartilage, as was done in Eskelinen et al. [18], to obtain a spatial distribution of these factors, which are known to influence cartilage metabolism.

It is important to note that the intracellular regulatory network for chondrocyte mechanotransduction and inflammation was developed using a combined data-driven and knowledge-based approach. For the data-driven component, a set of microarray databases from studies on mice was used [38]. It is worth mentioning that a detailed database specifically for human cartilage did not exist in the literature. While developing the network, it was ensured that the pathways included in the final model were common across species. For calibration and preliminary validation, the experimental study by Li et al. [12] was considered, which used bovine cartilage explants. Since gene expression in that study was obtained using qPCR experiments, no spatial data of gene expression was available for comparison with the corresponding numerical models. In the future, the use of spatial transcriptomics will provide spatial gene expression data of cartilage for a more detailed validation of the numerical multiscale model [49].

During validation of the multiscale model, ACAN gene expression was overpredicted relative to experimental measurements for both 10% and 20% dynamic compression of cartilage explants (Figure 5). This overprediction was accompanied by an underprediction of ADAMTS5, a major aggrecanase responsible for aggrecan degradation in cartilage [50]. It should be noted that in the experiments by Li et al. [12], gene expression was measured 48 hours after the onset of mechanical loading and cytokine treatment. In the multiscale model, gene expressions were recorded when the intracellular regulatory network reached a steady state numerically, without an explicit physical time scale. Given the inherently transient nature of gene expression, this discrepancy in temporal interpretation could likely contribute to the observed mismatch between model predictions and experimental data.

For further model validation, sGAG loss measured experimentally under different loading conditions was compared with the loss of aggrecan predicted by the multiscale model under comparable simulated conditions (Figure 5). In the model, the global activity of aggrecan was used as a surrogate measure for sGAG content, enabling a qualitative comparison with experimentally measured sGAG loss. This represents a simplified representation, as sGAG accumulation and release are influenced by post-translational modifications, extracellular processing, and transport phenomena not explicitly modeled in this study. Importantly, the model was not intended to predict absolute sGAG concentrations, but rather to assess the trends in matrix synthesis and loss under mechanical stimulation.

The next step in validation of the multiscale model will be to separately validate the individual length scales and their connections. To do so, in the future, multiscale experiments need to be performed that can measure both the mechanics at the tissue and cell scales simultaneously, as well as capture intracellular behavior by measuring the cellular response in terms of gene expressions and cell-secreted factors. Doing such experiments in a traditional bioreactor setup can be difficult. An alternative could be the use of microfluidic devices, such as those developed by Paggi et al. [51, 52] and Occhetta et al. [53], which can provide precise experimental data across multiple length scales. Another alternative could be the use of laser scanning microscopy approaches, as developed in [54–56], which enable real-time measurement of chondrocyte deformations during loading of cartilage explants.

Lastly, the multiscale modeling workflow developed in this study is unidirectional in nature, where the information flow is from the tissue scale to the intracellular scale via the cellular scale. In the future, efforts can be made to complete the feedback loop, i.e., to incorporate how changes in intracellular behavior can, in turn, affect the cellular microenvironment and thereby change the global mechanical behavior at the tissue scale.

In conclusion, the multiscale modeling workflow developed in this study is a first step towards a powerful tool for understanding the complex interplay of mechanics and inflammation in articular cartilage. By integrating tissue, cellular, and intracellular scale models, it provides a more comprehensive framework for studying cartilage mechanobiology. This approach facilitates a deeper understanding of the pathophysiology of OA and can provide valuable insights that can guide the development of targeted therapeutic strategies.

## Acknowledgements

This study received funding from the European Union’s Horizon 2020 research and innovation programme under the Marie Skłodowska-Curie grant agreement No. 721432 (CarBon project), the In Silico World project (grant agreement No. 101016503), the European Research Council Consolidator Grant No. 101088919 and the Belgian Federal Public Service Policy & Support (grant DigiTwin4PH). The funding sources have no role in design and execution of the study.

## References

[1] T. M. Griffin, F. Guilak, The role of mechanical loading in the onset and progression of osteoarthritis, Exerc Sport Sci Rev 33 (4) (2005) 195–200. doi:10.1097/00003677-200510000-00008.

[2] J. D. e. a. Steinmetz, Global, regional, and national burden of osteoarthritis, 1990–2020 and projections to 2050: a systematic analysis for the Global Burden of Disease Study 2021, The Lancet Rheumatology 5 (9). doi:10.1016/S2665-9913(23)00163-7. URL https://www.sciencedirect.com/science/article/pii/S2665991323001637

[3] R. F. Loeser, Integrins and chondrocyte–matrix interactions in articular cartilage, Matrix Biology 39 (2014) 11–16. doi:10.1016/j.matbio.2014.08.007. URL https://www.sciencedirect.com/science/article/pii/S0945053X14001607

[4] C. J. O’Conor, H. A. Leddy, H. C. Benefield, W. B. Liedtke, F. Guilak, TRPV4-mediated mechanotransduction regulates the metabolic response of chondrocytes to dynamic loading, Proc Natl Acad Sci U S A 111 (4) (2014) 1316–1321. doi:10.1073/pnas.1319569111.

[5] W. Lee, H. A. Leddy, Y. Chen, S. H. Lee, N. A. Zelenski, A. L. McNulty, J. Wu, K. N. Beicker, J. Coles, S. Zauscher, J. Grandl, F. Sachs, F. Guilak, W. B. Liedtke, Synergy between Piezo1 and Piezo2 channels confers high-strain mechanosensitivity to articular cartilage, Proc. Natl. Acad. Sci. U.S.A. 111 (47) (Nov. 2014). doi:10.1073/pnas.1414298111. URL https://pnas.org/doi/full/10.1073/pnas.1414298111

[6] A. P. Hollander, I. Pidoux, A. Reiner, C. Rorabeck, R. Bourne, A. R. Poole, Damage to type II collagen in aging and osteoarthritis starts at the articular surface, originates around chondrocytes, and extends into the cartilage with progressive degeneration, J Clin Invest 96 (6) (1995) 2859–2869. doi:10.1172/JCI118357.

[7] J. H. Rosenberg, V. Rai, M. F. Dilisio, D. K. Agrawal, Damage Associated Molecular Patterns in the Pathogenesis of Osteoarthritis: Potentially Novel Therapeutic Targets, Mol Cell Biochem 434 (1-2) (2017) 171–179. doi:10.1007/s11010-017-3047-4. URL https://www.ncbi.nlm.nih.gov/pmc/articles/PMC5671379/

[8] Z. Fan, S. Söder, S. Oehler, K. Fundel, T. Aigner, Activation of Interleukin-1 Signaling Cascades in Normal and Osteoarthritic Articular Cartilage, Am J Pathol 171 (3) (2007) 938–946. doi:10.2353/ajpath.2007.061083. URL https://www.ncbi.nlm.nih.gov/pmc/articles/PMC1959501/

[9] M. Neidlin, S. Dimitrakopoulou, L. G. Alexopoulos, Multi-tissue network analysis for drug prioritization in knee osteoarthritis, Sci Rep 9 (1) (2019) 15176. doi:10.1038/s41598-019-51627-6. URL https://www.nature.com/articles/s41598-019-51627-6

[10] M. Kapoor, J. Martel-Pelletier, D. Lajeunesse, J.-P. Pelletier, H. Fahmi, Role of proinflammatory cytokines in the pathophysiology of osteoarthritis, Nat Rev Rheumatol 7 (1) (2011) 33–42. doi:10.1038/nrrheum.2010.196. URL https://www.nature.com/articles/nrrheum.2010.196

[11] J. Nam, B. D. Aguda, B. Rath, S. Agarwal, Biomechanical Thresholds Regulate Inflammation through the NF-kB Pathway: Experiments and Modeling, PLoS One 4 (4) (2009) e5262. doi:10.1371/journal.pone.0005262. URL https://www.ncbi.nlm.nih.gov/pmc/articles/PMC2667254/

[12] Y. Li, E. H. Frank, Y. Wang, S. Chubinskaya, H.-H. Huang, A. J. Grodzinsky, Moderate dynamic compression inhibits pro-catabolic response of cartilage to mechanical injury, tumor necrosis factor-a and interleukin-6, but accentuates degradation above a strain threshold, Osteoarthritis and Cartilage 21 (12) (2013) 1933–1941. doi:10.1016/j.joca.2013.08.021. URL https://www.oarsijournal.com/article/S1063-4584(13)00939-4/fulltext

[13] V. D. Sree, A. B. Tepole, Computational systems mechanobiology of growth and remodeling: Integration of tissue mechanics and cell regulatory network dynamics, Current Opinion in Biomedical Engineering 15 (2020) 75–80. doi:10.1016/j.cobme.2020.01.002. URL https://www.sciencedirect.com/science/article/pii/S2468451120300039

[14] S. Mukherjee, M. Nazemi, I. Jonkers, L. Geris, Use of Computational Modeling to Study Joint Degeneration: A Review, Front. Bioeng. Biotechnol. 8 (Feb. 2020). doi:10.3389/fbioe.2020.00093. URL https://www.frontiersin.org/journals/bioengineering-and-biotechnology/articles/10.3389/fbioe.2020.00093/full

[15] V. B. Shim, P. J. Hunter, P. Pivonka, J. W. Fernandez, A Multiscale Framework Based on the Physiome Markup Languages for Exploring the Initiation of Osteoarthritis at the Bone–Cartilage Interface, IEEE Transactions on Biomedical Engineering 58 (12) (2011) 3532–3536. doi:10.1109/TBME.2011.2165955. URL https://ieeexplore.ieee.org/document/5997300

[16] G. A. Orozco, P. Bolcos, A. Mohammadi, M. S. Tanaka, M. Yang, T. M. Link, B. Ma, X. Li, P. Tanska, R. K. Korhonen, Prediction of local fixed charge density loss in cartilage following ACL injury and reconstruction: A computational proof-of-concept study with MRI follow-up, Journal of Orthopaedic Research 39 (5) (2021) 1064–1081. doi:10.1002/jor.24797. URL https://onlinelibrary.wiley.com/doi/abs/10.1002/jor.24797

[17] A. S. A. Eskelinen, M. E. Mononen, M. S. Venäläinen, R. K. Korhonen, P. Tanska, Maximum shear strain-based algorithm can predict proteoglycan loss in damaged articular cartilage, Biomechanics and Modeling in Mechanobiology 18 (3) (2019) 753–778. doi:10.1007/s10237-018-01113-1. URL https://doi.org/10.1007/s10237-018-01113-1

[18] A. S. A. Eskelinen, P. Tanska, C. Florea, G. A. Orozco, P. Julkunen, A. J. Grodzinsky, R. K. Korhonen, Mechanobiological model for simulation of injured cartilage degradation via pro-inflammatory cytokines and mechanical stimulus, PLoS Comput Biol 16 (6) (2020) e1007998. doi:10.1371/journal.pcbi.1007998.

[19] M. Segarra-Queralt, K. Crump, A. Pascuet-Fontanet, B. Gantenbein, J. Noailly, The interplay between biochemical mediators and mechanotransduction in chondrocytes: Unravelling the differential responses in primary knee osteoarthritis, Physics of Life Reviews 48 (2024) 205–221. doi:10.1016/j.plrev.2024.02.003. URL https://www.sciencedirect.com/science/article/pii/S1571064524000101

[20] M. Segarra-Queralt, G. Piella, J. Noailly, Network-based modelling of mechano-inflammatory chondrocyte regulation in early osteoarthritis, Front. Bioeng. Biotechnol. 11 (Feb. 2023). doi:10.3389/fbioe.2023.1006066. URL https://www.frontiersin.org/journals/bioengineering-and-biotechnology/articles/10.3389/fbioe.2023.1006066/full

[21] S. Khurana, S. Schivo, J. R. M. Plass, N. Mersinis, J. Scholma, J. Kerkhofs, L. Zhong, J. van de Pol, R. Langerak, L. Geris, M. Karperien, J. N. Post, An ECHO of Cartilage: In Silico Prediction of Combinatorial Treatments to Switch Between Transient and Permanent Cartilage Phenotypes With Ex Vivo Validation, Front Bioeng Biotechnol 9 (2021) 732917. doi:10.3389/fbioe.2021.732917. URL https://www.ncbi.nlm.nih.gov/pmc/articles/PMC8634894/

[22] P. Tanska, M. E. Mononen, R. K. Korhonen, A multi-scale finite element model for investigation of chondrocyte mechanics in normal and medial meniscectomy human knee joint during walking, Journal of Biomechanics 48 (8) (2015) 1397–1406. doi:10.1016/j.jbiomech.2015.02.043. URL https://linkinghub.elsevier.com/retrieve/pii/S0021929015001335

[23] F. Guilak, W. R. Jones, H. Ting-Beall, G. M. Lee, The deformation behavior and mechanical properties of chondrocytes in articular cartilage, Osteoarthritis and Cartilage 7 (1) (1999) 59–70. doi:10.1053/joca.1998.0162. URL https://linkinghub.elsevier.com/retrieve/pii/S1063458498901625

[24] W. Wilson, C. C. van Donkelaar, B. van Rietbergen, K. Ito, R. Huiskes, Stresses in the local collagen network of articular cartilage: a poroviscoelastic fibril-reinforced finite element study, Journal of Biomechanics 37 (3) (2004) 357–366. doi:10.1016/S0021-9290(03)00267-7. URL https://www.sciencedirect.com/science/article/pii/S0021929003002677

[25] J. M. P. Quiroga, W. Wilson, K. Ito, C. C. van Donkelaar, Relative contribution of articular cartilage’s constitutive components to load support depending on strain rate, Biomech Model Mechanobiol 16 (1) (2017) 151–158. doi:10.1007/s10237-016-0807-0. URL https://doi.org/10.1007/s10237-016-0807-0

[26] S. Mukherjee, W. Wilson, L. Geris, From human joints to bioreactor setups: quantifying mechanical stimuli in cartilage physiology and regeneration, iSSN: 2692-8205 Pages: 2025.10.06.678667 Section: New Results (Oct. 2025). doi:10.1101/2025.10.06.678667. URL https://www.biorxiv.org/content/10.1101/2025.10.06.678667v1

[27] A. Heuijerjans, W. Wilson, K. Ito, C. C. Van Donkelaar, Osteochondral resurfacing implantation angle is more important than implant material stiffness, Journal Orthopaedic Research 36 (11) (2018) 2911–2922. doi:10.1002/jor.24101. URL https://onlinelibrary.wiley.com/doi/10.1002/jor.24101

[28] P. Julkunen, W. Wilson, J. S. Jurvelin, R. K. Korhonen, Composition of the pericellular matrix modulates the deformation behaviour of chondrocytes in articular cartilage under static loading., Medical & biological engineering & computing 47 (12) (2009) 1281–90. doi:10.1007/s11517-009-0547-8. URL http://www.ncbi.nlm.nih.gov/pubmed/19898885http://www.pubmedcentral.nih.gov/articlerender.fcgi?artid=PMC2779377

[29] S. C. Sibole, A. Erdemir, Chondrocyte Deformations as a Function of Tibiofemoral Joint Loading Predicted by a Generalized High-Throughput Pipeline of Multi-Scale Simulations, PLoS ONE 7 (5) (2012) e37538. doi:10.1371/journal.pone.0037538. URL http://dx.plos.org/10.1371/journal.pone.0037538

[30] S. C. Sibole, S. Maas, J. P. Halloran, J. A. Weiss, A. Erdemir, Evaluation of a post-processing approach for multiscale analysis of biphasic mechanics of chondrocytes, Computer Methods in Biomechanics and Biomedical Engineering 16 (10) (2013) 1112–1126. doi:10.1080/10255842.2013.809711. URL https://doi.org/10.1080/10255842.2013.809711

[31] M. Kääb, R. Richards, K. Ito, I. ap Gwynn, H. Nötzli, Deformation of Chondrocytes in Articular Cartilage under Compressive Load: A Morphological Study, Cells Tissues Organs 175 (3) (2003) 133–139. doi:10.1159/000074629. URL https://doi.org/10.1159/000074629

[32] C. A. POOLE, Review. Articular cartilage chondrons: form, function and failure, J Anat 191 (Pt 1) (1997) 1–13. doi:10.1046/j.1469-7580.1997.19110001.x. URL https://www.ncbi.nlm.nih.gov/pmc/articles/PMC1467653/

[33] C. A. Poole, S. Ayad, R. T. Gilbert, Chondrons from articular cartilage: V.* Immunohistochemical evaluation of type VI collagen organisation in isolated chondrons by light, confocal and electron microscopy, Journal of Cell Science 103 (4) (1992) 1101–1110. doi:10.1242/jcs.103.4.1101. URL https://doi.org/10.1242/jcs.103.4.1

[34] I. Youn, J. Choi, L. Cao, L. Setton, F. Guilak, Zonal variations in the three-dimensional morphology of the chondron measured in situ using confocal microscopy, Osteoarthritis and Cartilage 14 (9) (2006) 889–897. doi:10.1016/j.joca.2006.02.017. URL https://linkinghub.elsevier.com/retrieve/pii/S1063458406000471

[35] H. Guo, P. A. Torzilli, Shape of chondrocytes within articular cartilage affects the solid but not the fluid microenvironment under unconfined compression, Acta Biomater 29 (2016) 170–179. doi:10.1016/j.actbio.2015.10.035. URL https://www.ncbi.nlm.nih.gov/pmc/articles/PMC4681666/

[36] J. Z. Kechagia, J. Ivaska, P. Roca-Cusachs, Integrins as biomechanical sensors of the microenvironment, Nat Rev Mol Cell Biol 20 (8) (2019) 457–473. doi:10.1038/s41580-019-0134-2. URL https://www.nature.com/articles/s41580-019-0134-2

[37] A. Saraswathibhatla, D. Indana, O. Chaudhuri, Cell-extracellular matrix mechanotransduction in 3D, Nat Rev Mol Cell Biol 24 (7) (2023) 495–516. doi:10.1038/s41580-023-00583-1.

[38] R. Lesage, M. N. Ferrao Blanco, R. Narcisi, T. Welting, G. J. V. M. van Osch, L. Geris, An integrated in silico-in vitro approach for identifying therapeutic targets against osteoarthritis, BMC Biol 20 (1) (2022) 253. doi:10.1186/s12915-022-01451-8. URL https://doi.org/10.1186/s12915-022-01451-8

[39] A. Mobasheri, C. Matta, I. Uzielienè, E. Budd, P. Martín-Vasallo, E. Bernotiene, The chondrocyte channelome: A narrative review, Joint Bone Spine 86 (1) (2019) 29–35. doi:10.1016/j.jbspin.2018.01.012.

[40] R. J. Nims, L. Pferdehirt, F. Guilak, Mechanogenetics: harnessing mechanobiology for cellular engineering, Current Opinion in Biotechnology 73 (2022) 374–379. doi:10.1016/j.copbio.2021.09.011. URL https://linkinghub.elsevier.com/retrieve/pii/S0958166921001828

[41] S. Madhavan, M. Anghelina, D. Sjostrom, A. Dossumbekova, D. C. Guttridge, S. Agarwal, Biomechanical Signals Suppress TAK1 Activation to Inhibit NF-kB Transcriptional Activation in Fibrochondrocytes1, The Journal of Immunology 179 (9) (2007) 6246–6254. doi:10.4049/jimmunol.179.9.6246. URL https://doi.org/10.4049/jimmunol.179.9.6246

[42] J. Kerkhofs, L. Geris, A Semiquantitative Framework for Gene Regulatory Networks: Increasing the Time and Quantitative Resolution of Boolean Networks, PLOS ONE 10 (6) (2015) e0130033. doi:10.1371/journal.pone.0130033. URL https://dx.plos.org/10.1371/journal.pone.0130033

[43] T. Hodgkinson, D. C. Kelly, C. M. Curtin, F. J. O’Brien, Mechanosignalling in cartilage: an emerging target for the treatment of osteoarthritis, Nat Rev Rheumatol 18 (2) (2022) 67–84. doi:10.1038/s41584-021-00724-w. URL https://www.nature.com/articles/s41584-021-00724-w

[44] Z. Karagöz, L. Rijns, P. Y. W. Dankers, M. van Griensven, A. Carlier, Towards understanding the messengers of extracellular space: Computational models of outside-in integrin reaction networks, Computational and Structural Biotechnology Journal 19 (2021) 303–314. doi:10.1016/j.csbj.2020.12.025. URL https://www.sciencedirect.com/science/article/pii/S200103702030550X

[45] M. Almonte-Becerril, M. Costell, J. B. Kouri, Changes in the integrins expression are related with the osteoarthritis severity in an experimental animal model in rats, Journal of Orthopaedic Research 32 (9) (2014) 1161–1166. doi:10.1002/jor.22649. URL https://onlinelibrary.wiley.com/doi/abs/10.1002/jor.22649

[46] W. Lee, R. J. Nims, A. Savadipour, Q. Zhang, H. A. Leddy, F. Liu, A. L. McNulty, Y. Chen, F. Guilak, W. B. Liedtke, Inflammatory signaling sensitizes Piezo1 mechanotransduction in articular chondrocytes as a pathogenic feed-forward mechanism in osteoarthritis, Proc. Natl. Acad. Sci. U.S.A. 118 (13) (2021) e2001611118. doi:10.1073/pnas.2001611118. URL https://pnas.org/doi/full/10.1073/pnas.2001611118

[47] W. Gao, H. Hasan, D. E. Anderson, W. Lee, The Role of Mechanically-Activated Ion Channels Piezo1, Piezo2, and TRPV4 in Chondrocyte Mechanotransduction and Mechano-Therapeutics for Osteoarthritis, Front. Cell Dev. Biol. 10 (2022) 885224. doi:10.3389/fcell.2022.885224. URL https://www.frontiersin.org/articles/10.3389/fcell.2022.885224/full

[48] A. Savadipour, R. J. Nims, N. Rashidi, J. M. Garcia-Castorena, R. Tang, G. K. Marushack, S. J. Oswald, W. B. Liedtke, F. Guilak, Membrane stretch as the mechanism of activation of PIEZO1 ion channels in chondrocytes, Proc. Natl. Acad. Sci. U.S.A. 120 (30) (2023) e2221958120. doi:10.1073/pnas.2221958120. URL https://pnas.org/doi/10.1073/pnas.2221958120

[49] X. Fan, A. R. Sun, R. S. E. Young, I. O. Afara, B. R. Hamilton, L. J. Y. Ong, R. Crawford, I. Prasadam, Spatial analysis of the osteoarthritis microenvironment: techniques, insights, and applications, Bone Res 12 (1) (2024) 1–19. doi:10.1038/s41413-023-00304-6. URL https://www.nature.com/articles/s41413-023-00304-6

[50] H. Stanton, F. M. Rogerson, C. J. East, S. B. Golub, K. E. Lawlor, C. T. Meeker, C. B. Little, K. Last, P. J. Farmer, I. K. Campbell, A. M. Fourie, A. J. Fosang, ADAMTS5 is the major aggrecanase in mouse cartilage in vivo and in vitro, Nature 434 (7033) (2005) 648–652. doi:10.1038/nature03417.

[51] C. A. Paggi, B. Venzac, M. Karperien, J. C. H. Leijten, S. Le Gac, Monolithic microfluidic platform for exerting gradients of compression on cell-laden hydrogels, and application to a model of the articular cartilage, Sensors and Actuators B: Chemical 315 (2020) 127917. doi:10.1016/j.snb.2020.127917. URL https://www.sciencedirect.com/science/article/pii/S0925400520302653

[52] C. A. Paggi, L. M. Teixeira, S. Le Gac, M. Karperien, Joint-on-chip platforms: entering a new era of in vitro models for arthritis, Nat Rev Rheumatol 18 (4) (2022) 217–231. doi:10.1038/s41584-021-00736-6. URL https://www.nature.com/articles/s41584-021-00736-6

[53] P. Occhetta, A. Mainardi, E. Votta, Q. Vallmajo-Martin, M. Ehrbar, I. Martin, A. Barbero, M. Rasponi, Hyperphysiological compression of articular cartilage induces an osteoarthritic phenotype in a cartilage-on-a-chip model, Nat Biomed Eng 3 (7) (2019) 545–557. doi:10.1038/s41551-019-0406-3. URL https://www.nature.com/articles/s41551-019-0406-3

[54] A. Komeili, B. S. Otoo, Z. Abusara, S. Sibole, S. Federico, W. Herzog, Chondrocyte Deformations Under Mild Dynamic Loading Conditions, Ann Biomed Eng 49 (2) (2021) 846–857. doi:10.1007/s10439-020-02615-9. URL https://doi.org/10.1007/s10439-020-02615-9

[55] S. C. Sibole, E. K. Moo, S. Federico, W. Herzog, Dynamic Deformation Calculation of Articular Cartilage and Cells Using Resonance-Driven Laser Scanning Microscopy, Journal of Biomechanical Engineering 145 (021005) (Oct. 2022). doi:10.1115/1.4055308. URL https://doi.org/10.1115/1.4055308

[56] B. S. Otoo, E. Kuan Moo, A. Komeili, D. A. Hart, W. Herzog, Chondrocyte deformation during the unloading phase of cyclic compression loading, Journal of Biomechanics 171 (2024) 112179. doi:10.1016/j.jbiomech.2024.112179. URL https://www.sciencedirect.com/science/article/pii/S0021929024002574

