## Supplementary Materials for "A multiscale modeling approach to study the role of mechanics and inflammation in pathophysiology of articular cartilage"

---

### S1. Detailed explanation of the pathways involved in the regulatory network

This supplementary data contains detailed explanations of the different pathways included in the regulatory networks (adapted from Lesage et al. 2021)

#### *S1.1. Mechanotransduction pathways*

##### *S1.1.1. Integrin mechanoreceptor and downstream pathways*

One of the major sensor by which chondrocyte sense the external forces are the transmembrane integrin proteins and focal adhesion complexes. These complexes are made of integrins and other cytoplasmic molecules, which provides kinase activity to the complexes (14, 15). They mediate mechanical force transmission to downstream biochemical reaction cascades. There are several integrin isoforms and subunits that differs between species (e.g. $\alpha1\beta1$, $\alpha5\beta1$, etc..) and that have binding sites for different ECM fibers (e.g. collagen, fibronectin), which act as external ligands (16). The $\alpha5\beta1$subunint has been shown to be important for modulating the expression of MMPs in mouse chondrocytes (17), but too few studies distinguish downstream cascades and effects specific to each integrin subunits. Thus, the node representing integrins in the model is a general summarization of a receptor translating mechanical cues into biochemical cascades, for sake of simplification. One specific human gene coding for this type of protein could be ITGB1, for instance. Generally speaking, focal adhesion kinases (FAK) are among of the main effectors of focal adhesion complexes, for signal transduction (18). More actors are involved to offer refined response to various loading regimen but, for sake of simplification, our model summarizes these molecular complexes by a single node named FAK. This type of kinases functions as a hub to transduce signals, including through phosphorylation of the RhoA GTPases, which are known to be involved in the regulation of cell response to mechanical cues (19). More particularly, these phosphorylation cascades are involved in the regulation of cell differentiation by mechanical cues in development and regenerative processes (20). RhoA proteins and the downstream ROCK1/2 proteins regulate the actin cytoskeleton assembly and contractility by phosphorylating myosin light chains (MLC) (21). This level of contractility and cytoskeleton assembly is represented by a single variable named Actin-MLC in the model. Evidences showed that RhoA and ROCK are involved in the suppression of hypertrophic chondrocyte differentiation (22), possibly in part through activation of the cyclin CCND1, a known suppressor of RUNX2 activity (22), which is accounted for in the model. Actin-myosin phosphorylation by the RhoA/ROCK axis also takes place through LIM-domain containing protein kinase and cofilin proteins (21, 23), but this side and redundant pathway that does not involve many crosstalk with other actors was not represented in the model for sake of simplification. Other downstream effectors below the RhoA/ROCK axis are multiple. First, ROCK would promote SOX9 phosphorylation, thereby promoting its transduction to nucleus in chondrocyte (24). SOX9 is commonly known to directly upregulate expression of the genes producing ECM proteins such as proteoglycans and type-II collagen (24). In other words, this axis can fairly be presumed as a mechanically induced pro-anabolic pathway, when considered in isolation of the full connectome. Second, the phosphorylation of the cytoskeleton stress fibers and MLC, which increases the stability and contractility of the cytoskeleton, regulates the transcriptional activity of various TFs (25). For instance, it promotes the nuclear translocation of $\beta$-catenin, an effector of the Wnt canonical pathway, thereby promoting the WNT/$\beta$-catenin axis (20). In the same way, it also promotes the activation of Yes-associated protein (YAP) and the transcriptional co-activator with PDZ-binding motif (TAZ). Actin contractility would also up-regulate ECM related genes such as COL2A1 and ACAN (or Aggrecan, a proteoglycan), but the exact nature of the involved TF is not clear. For that reason, we introduced a dummy TF in the model, for the transcriptional activity induced by the cytoskeleton-increased contractility (more details below). Mechanical stimulation of chondrocytes increased their sensitivity to mechanical cues, suggesting the involvement of a positive feedback loop. There are experimental evidences that cyclic loading increases integrin receptors amount at the membrane after mechanical stimulation (26). We propose a mechanism in which, upon integrin stimulation, downstream pathways promote the integrin gene up-regulation, thereby increasing the abundance of integrin receptors. In the model, we assume that this positive transcriptional activity happens through the dummy-TF introduced earlier and lying downstream of the RhoA/ROCK/Actin-MLC pathway, the main force-transducing axis. A possible candidate for that dummy-TF could be serum response factor (SRF) because the Rho family of GTPase (RhoA, Rac1) as well as actin contractility are reported to promote the SRF transcription factor activation upon mechanical stimulation and that factor would regulate cell contractility related genes in return (15, 27, 28). In absence of specific information, this dummy-TF or SRF is not regulated at the transcriptional level, in the model. As mentioned previously, YAP and TAZ have been identified to be mechanosensitive and to be involved in the transduction of mechanical signals (29). They can be activated or deactivated according to the mechanical environment, such as ECM stiffness. YAP and TAZ are considered as analogous; nevertheless, downstream effects are often reported for YAP rather than TAZ, in literature (30). In the current model, we do not make any distinction between the two and they are represented by a single entity. YAP inactivation is induced by phosphorylation, which prevent its translocation to the nucleus and may even direct it toward proteasome degradation (31). Mechanically-induced YAP/TAZ activation, downstream of the RhoA/ROCK/Actin-MLC axis, would be independent from the often reported NF2/Hippo/LATS biochemical pathway (29). For that reason the Hippo pathway was not represented in the model. Furthermore, YAP and TAZ are involved in the regulation of chondrogenic phenotype in response to physical cues (20, 30, 32). For instance, YAP was reported to block hypertrophy during skeletal development (30) while the overall amount of YAP is reduced during OA (31). Mechanistically, YAP contributes to SOX6 upregulation and reduces RUNX2 activity by direct binding, eventually preventing RUNX2 to upregulate COL10A1 expression (30). Our mathematical model accounts for these mechanisms by letting YAP/TAZ increase SOX9 at genetic level, which represents the SOX trio (SOX9, 5 and 6), and decrease RUNX2 protein activation. With the lack of specific information about their transcriptional regulators in cartilage, YAP/TAZ are only modulated at the protein level and not at the transcriptional level in the present numerical model. Finally, mTOR is a well-known mechanosensitive gene that is activated by cyclic loading in chondrocytes (33). This mechanically induced activation was reported to happen downstream of the PI3K/AKT/mTOR cascade in osteoblasts. This axis being also active in chondrocytes we introduced an activation link directly from integrin to PI3K, in the model. Inhibition of mTOR have been shown to reduce OA severity (34) but the implications of mTOR in chondrocyte biology are manifold and we only introduced a few downstream effectors, in the model. We modeled mTOR capacity to phosphorylate and help the nuclear translocation of GLI2 (35), the nuclear effector the IHH pathway. This interaction also accounts for the ability of mTOR to regulate the PTHrP pathway via GLI2 in chondrocytes (35) since GLI2 was already modelled as an upstream positive regulator of PTHrP expression. In addition, mTOR may enhance RUNX2 expression through indirect activation of the transcription factor DLX5 and GLI2 (36) and is capable of suppressing its own inhibitor PP2A in the model (37, 38), generating a positive feedback loop. The involvement of mTOR in different complexes (i.e. mTORC1 and mTORC2) was not considered and many mTOR-related mechanisms were disregarded in the current model.

*S1.1.2. Ion channel and calcium signaling*

Beside integrins, other important mechanosensors are expressed by chondrocytes, including the the so called chondrocyte “channelome”. We focused on the cations channels entitled transient receptor potential vanilloid 4 (TRPV4), largely located at the cell membrane of articular chondrocyte (39). They are multimodally activated channels as they can transduce dynamic compressive loading signals and be activated by osmotic pressure variations as well as chemical signals (40, 41). TRPV4 plays a major role in the physiological response to cartilage loading by mediating intracellular calcium cations (Ca2+) influx (40). Functional mutation of the TRPV4 channel have been related to skeletal dysplasia and joint degenerative conditions such as osteoarthritis in human (40, 42). TRPV4 expression and activation was often related to SOX9 increased expression as well as overexpression of ECM related proteins such as type II-collagen and proteoglycans, as such, it is believed to be involved in chondrogenic differentiation (43). Importantly, Ca2+ signaling is an essential universal secondary messenger in articular chondrocytes and its regulation is reported to be highly complex and based on frequency of concentration oscillations (44, 45). The current model being discrete in time and semi-quantitative, it does not allow to represent this tightly regulated periodic flux, which could be subject to a dedicated time continuous model in itself. Instead, in our model, the TRPV4 entity can be activated with increased mechanical loading, based on the same forces that are sensed by integrins, it subsequently increases the activity of the node called Ca2+ representing the concentration or effective level of cytoplasmic Ca2+. In that sense, we can consider that the maximal value of the variable would correspond to the most optimal level and frequency of Ca2+ oscillations and the lower value to less physiological frequencies, since the spontaneous calcium signaling is known to be altered in disease (46). Actually, both intra (e.g. endoplasmic reticulum) and extra-cellular sources of Ca2+ exist but the current model focuses on the extracellular source of influx as supported by this study (44) according to which Ca2+ signaling, either spontaneous or load induced, has to be initiated by the extracellular Ca2+ influx. cAMP response element-binding protein (CREB) is one of the downstream transcription factors which decipher the information encoded by Ca2+

concentration, this connection was represented by an activation link toward ATF4 (also known as CREB2), which was already present in the regulatory model. TRPV4 stimulation with dynamical mechanical loading or agonist treatment was reported to modulate ECM proteins biosynthesis. In particular, it increases COL2A1 expression and decreases the mRNA level of ADAMTS5, an ECM proteinase (40). The first observation was not directly encoded in the model but it is expected to be mediated by the activating effect of Ca2+ on the cytoskeleton contractility and on SOX9 expression in the model, as mentioned above. The second one is represented by an inhibitory edge relating Ca2+ activity and ADAMTS5 gene expression, even though that relationship is likely to be indirect and to involve an intermediary TF, which is not precisely identified, to our knowledge. Interestingly, this TRPV4-dependant mechanosensitive pathway can be modulated by, and interplay with, other pathways. For instance, N. Trompeter et al. have recently showed that the TRPV4 channel was regulated by the actin cytoskeleton through binding of F-actin to the Microtubule Associated Protein Binding 7 domain of TRPV4. They have also shown that TRPV4 activity is suppressed by IGF-1 treatment concluding that IGF-1 would modulate the activity of this channel through alteration in the actin cytoskeletal organization (47). Indeed, IGF-1 treatment in primary chondrocytes increased cell stiffness and F-actin formation (48, 49). In order to integrate this precise mechanism in the model, we made sure that concomitant activity of the IGF-1 receptor and F-Actin was able to decrease the TRPV4 protein activity (i.e. channel opening) by using a multiplicative factor in the equations. That way, the Actin-MLC can inhibit TRPV4 activity only if the IGF-1 receptor is activated by its ligand, and this in a dose dependent way. Moreover, this calcium signaling interacts with the Rho GTPase/Actin axis to promote F-actin stability and contractility (50), which is translated by an activation link between Ca2+ and RhoA/ROCK, in our model.

#### *S1.2. Inflammatory pathways*

Increasing evidences have highlighted the implications inflammatory events in OA onset and progression (51–53). Even though a relatively low synovitis grade is observed during OA, this pathology is characterized an auto-activating inflammatory loop. Indeed, in presence of some ECM degradation products or damage-associated molecular patterns (DAMPS), inflammatory receptors such as the Toll-like receptors trigger signal transductions resulting in pro-inflammatory cytokine production by the chondrocytes, themselves (3, 54–56). In this section, we summarize the main inflammatory related pathways that we integrated in our model and the assumptions made. First, several pro-inflammatory mediators, for which chondrocytes expressed the specific receptors (e.g. Toll-like receptor), are responsible for inducing pro-inflammatory events. For the sake of simplification in the model, we represented a single pro-inflammatory cytokine entity (e.g. Interleukine-1 beta (IL1B) or Tumor Necrosis Factor (TNF)) that is able to signal through a unique receptor of inflammation (R-inflam). This is an important assumption allowing us to focus on the intracellular interplay between mechanotransduction and inflammation rather than the role of specific receptors and signaling molecules. In chondrocytes, IL1B increases the production of proteases responsible for matrix degeneration, suppresses matrix biosynthesis and induces pro-inflammatory mediators by controlling expression of genes such as inducible NO synthase (iNOS) or matrix metalloproteinases (MMPs) (51, 57). In addition, pro-inflammatory cytokines like IL1B promote their own production, which amplifies the immune response as a positive feedback loop (58), This may be mediated by several signaling cascades involving TAK1, NFKB, and MAPKs, we refer the reader to Chapter 1 for more details and we shortly recall the main axes in the paragraphs below. In particular, TAK1 is a critical point where signals generated by pro-inflammatory signals converge, notably to initiate NFKB signaling cascade and pro-inflammatory gene induction (58, 59). The canonical mechanism of action involves a double inhibition and the release of NFKB from its inhibitor, IKBA (56). Since the other upstream activators of NFKB are only able to activate it for nuclear translocation when it is free from IKBA, we devised a special term in which the sum of the upstream activators is multiplied by the term (1- [level of IKBA]) in the model's equation. In addition, there exists a negative feedback loop in which NFKB promotes the gene expression of its own inhibitor IKBA(56). iNOS is an enzyme that can be induced by pro-inflammatory cytokines and that promotes the production of nitrite oxide, a pro-inflammatory mediator feeding the inflammatory loop in abnormal situations (60, 61). Indeed, in cytokine-stimulated chondrocytes, induced-nitric oxide causes persistent activation of NFKB (60). The inflammation inducible effect was represented in the model by overexpression of iNOS by NFKB, since the promoter region of the iNOS gene contains an NFKB binding site (60, 62), and the loop feeding with the direct activation of NFKB by NO. Another path involved in the activation of chondrocyte inflammatory response is the Wnt signaling. Indeed, overexpression of TCF4, a WNT downstream effector, was shown to induce the expression of MMPs and the activation of NFKB in chondrocytes. TCF would directly bind to NFKB to help its nuclear translocation in articular chondrocytes (63). This was accounted for by a direct activation link between LEF/TCF and NFKB at the protein level. Nevertheless, the aforementioned auto-inflammation loop does not function in a fully uncontrolled fashion. There exist suppressors of cytokine signaling (e.g. SOCS proteins), which are soluble inhibitors of pro-inflammatory cytokines receptors (Rinflam, in the model) (59). They prevent the downstream phosphorylation cascades that normally lead to NFKB activation and negatively regulate the cytokine induced-JAK/STAT pathway during osteoarthritis (64). SOCS proteins are upregulated by the FGF and IGFI pathways, which illustrates some of the crosstalk happening with other signaling pathways, in the model. Another way that IL1B-R mediated signaling can be terminated is by activating some negative feedback loops. For instance, the p38 MAPK induces the phosphorylation of TAB1, which inactivates TAK1, and NFKB can overexpresses its own inhibitor IKBA. Inflammatory signal also interplays with IGF signaling, in several ways. One way involves the IGF binding proteins, which bind to IGFI and prevent downstream signaling. These proteins are known to be up-regulated during hypertrophy and OA (65, 66). While IGFI signaling may act as a break to inflammatory signal via up-regulation of SOCS proteins, IL-1 promotes the production and release of IGFBPs (67), which is translated by the up-regulation of IGFBP expression by the nuclear effector NFKB in our model, although the real mode of action is not clear yet. Finally, FOXO is an important anti-catabolic TF modulated and downregulated by inflammation (68) and represented in the in silico model via its ability to promote proteoglycans and collagen production and downregulate MMP13 and ADAMTS5 (69).

*S1.3. Interplay between pathways*

Previously we compartmented pathways into the mechanoregulatory and inflammatory specific pathways but those cellular paths are actually interconnected. S. Madhavan et al. have showed that cyclic tensile strains suppressed pro-inflammatory gene induction induced by all the three inflammatory mediators, IL-1$\beta$, TNF-$\alpha$, and LPS (58). However, the exact relationship is complex and neither linear nor straightforward. Here we describe some of the most direct molecular mechanisms by which mechanical loading and inflammation influence each other and that are accounted for in the current computational model. First, signal transduction through integrin receptors such as integrin subunits $\alpha 5$ and $\beta 1$ have been shown to up-regulate MMP13 expression in chondrocyte, when associated with inflammatory cues. Indeed, that effect would likely be mediated through induction of IL1B (17, 26). It corroborates with the fact that excessive mechanical loading is thought to cause NFKB activation and OA development (10). However, this observation likely depends on the status of all other connected signaling pathways and the magnitude of the biomechanical signals since it was also repeatedly reported that mechanical signals at lower or physiological magnitudes maintain healthy cartilage and are potent inhibitors of inflammation and degrading enzymes (10, 58, 70). The TAK1/NFKB pathway has been suggested as key player of this biomechanical signal-mediated anti-inflammatory effect (59). In normal physiological ranges, mechanical stimulations would block TAK1 phosphorylation, thereby preventing IKK phosphorylation and activation (58, 59, 71). A possible mode of action that was integrated in the model regards YAP/TAZ. Indeed, YAP and NFKB pathways reciprocally inhibit each other, possibly affecting cartilage degradation in OA (31). On the one hand IL1B and TNFA promotes YAP/TAZ degradation through TAK1-mediated phosphorylation (31). On the other hand, YAP directly binds with TAK1, thereby limiting NFKB activation by blocking reaction substrate accessibility (31). Second, iNOS and NO are in a central position for the mechano-inflammatory crosstalk. Indeed, the pro-inflammatory mediator iNOs is also mechano-inducible (72, 73). This is represented in the model by a direct activatory link between Actin-MLC, the most direct intracellular mechanical transducer, and iNOS at protein level. However, the exact mode of action is not Third, ECM degradation products, also called damage-associated molecular patterns (DAMPs) are generated in case ADAMTS5 proteinases are released by chondrocytes or in case of adverse events, as observed in OA(74). Those degradation products interact with membrane receptors of cytokines such as Toll-like receptors (TLRs) but also integrins, eventually increasing pro-inflammatory and catabolic products. In the model we shortcut the implication of these DAMPs and we added a link describing the indirect activation of inflammatory cytokine receptors and integrins by ADAMTS5, assuming that ADAMTS5 amounts and DAMPS were linearly related for simplification. However, calcium signaling (TRPV4/Ca2+), which is mechanosensitive, inhibits ADAMTS5 expression in chondrocytes. This is another illustration of the multiple and complex connections between mechanotransduction and inflammatory responses. In the end, cells integrate all the aforementionned signals and modify their secretary profile accordingly, which affects their the matrix mechanical properties and cells mechanobiological environement (75), eventually leading to new external cues.

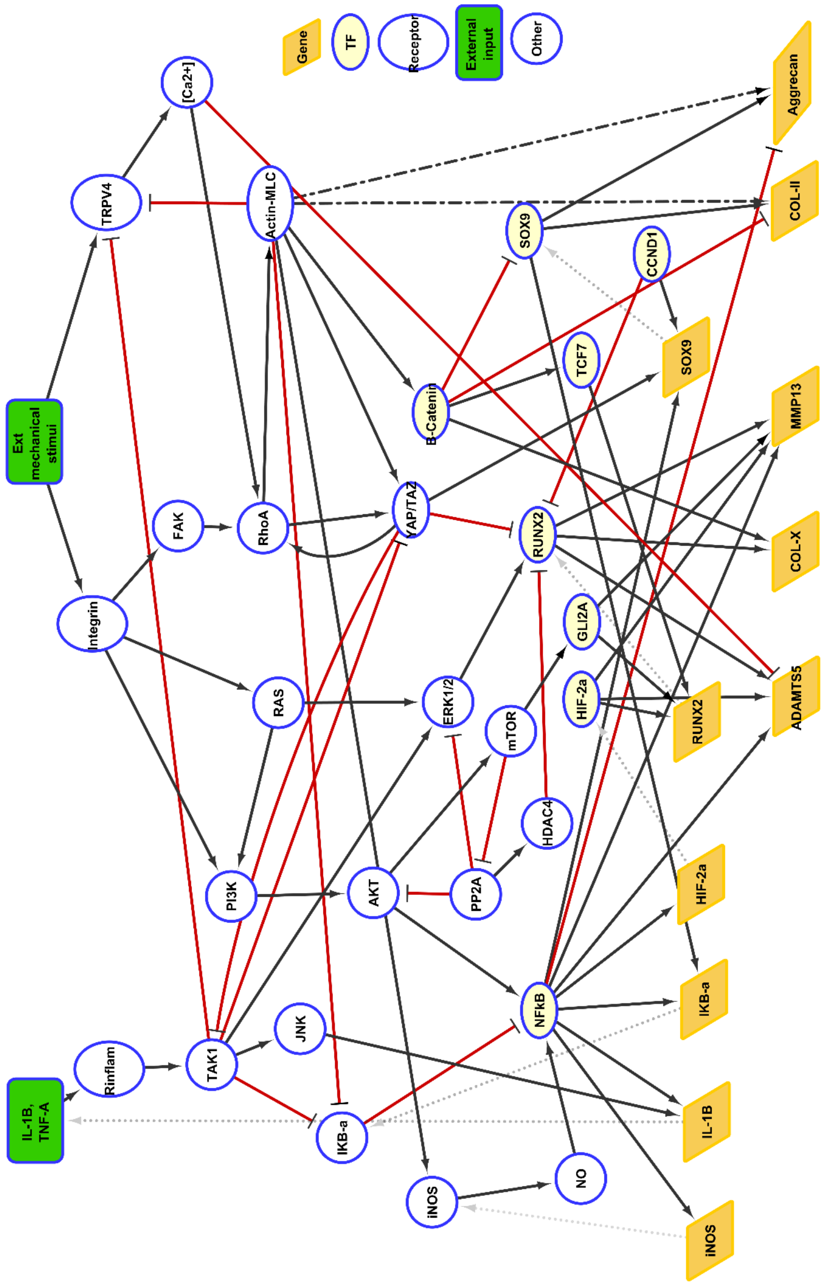


Figure S1: Snapshot of the mechanotransduction and pro-inflammatory signaling and their crosstalk in the in silico chondrocyte. This activity flow diagram is an arbitrary selection of important crosstalk from the model and related to mechanotransduction or inflammation, for illustration. External cues (in green) are either the mechanical loading or pro-inflammatory cytokines. The signal is transduced through signaling molecules ('Other') up to transcription factors ('TF' in blue circled yellow ellipse), which up- or down-regulate the expression of downstream genes (yellow parallelograms). As a result, genes increase the amount of the corresponding upstream proteins (grey dotted lines). Black arrows (resp. red T-ended lines) represent activation (resp. inhibition) interactions or influences. Influences may be direct or indirect in chondrocytes, the dashed black lines represent interactions that involve an intermediary node in the model but that is unrepresented in the graph.

### S2. Mesh convergence study of the cell level model

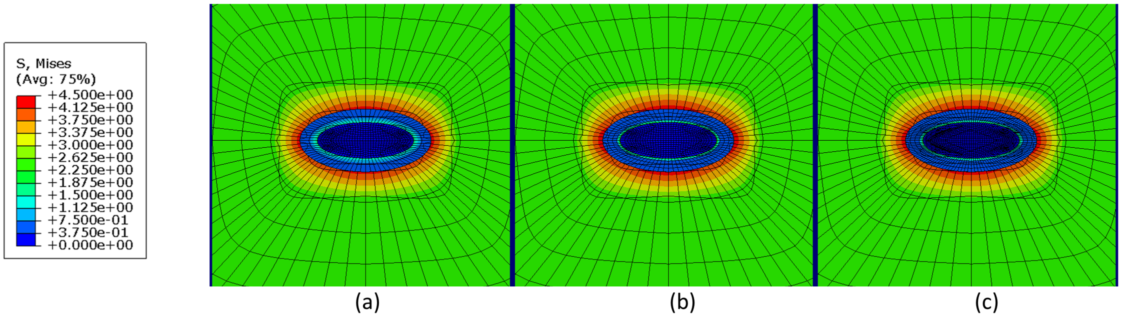


Figure S2: Mesh convergence study for cell-level model 10% strain. (a) Mesh=29216 elements. (b) Mesh=32912 elements. (c) Mesh=59092 elements.

**References**

1. A. C. Lin, B. L. Seeto, J. M. Bartoszko, M. A. Khoury, H. Whetstone, L. Ho, C. Hsu, A. S. Ali, B. A. Alman, Modulating hedgehog signaling can attenuate the severity of osteoarthritis. Nat. Med. 15, 1421–1425 (2009).

2. Z. Fan, S. Söder, S. Oehler, K. Fundel, T. Aigner, Activation of interleukin-1 signaling cascades in normal and osteoarthritis articular cartilage. Am. J. Pathol. 171, 938–946 (2007).

3. J. H. Rosenberg, V. Rai, M. F. Dilisio, D. K. Agrawal, Damage-associated molecular patterns in the pathogenesis of osteoarthritis: potentially novel therapeutic targets. Mol. Cell. Biochem. 434 (2017), pp. 171–179.

4. M. Neidlin, S. Dimitrakopoulou, L. G. Alexopoulos, Multi-tissue network analysis for drug prioritization in knee osteoarthritis. Sci. Rep. 9 (2019), doi:10.1038/s41598-019-51627-6.

5. M. Kapoor, J. Martel-Pelletier, D. Lajeunesse, J. P. Pelletier, H. Fahmi, Role of proinflammatory cytokines in the pathophysiology of osteoarthritis. Nat. Rev. Rheumatol. 7 (2011), pp. 33–42.

6. J. Nam, B. D. Aguda, B. Rath, S. Agarwal, Biomechanical Thresholds Regulate Inflammation through the NF-$\kappa$B Pathway: Experiments and Modeling. PLoS One. 4, e5262 (2009).

7. Y. Li, E. H. Frank, Y. Wang, S. Chubinskaya, H. H. Huang, A. J. Grodzinsky, Moderate dynamic compression inhibits pro-catabolic response of cartilage to mechanical injury, tumor necrosis factor-$\alpha$ and interleukin-6, but accentuates degradation above a strain threshold. Osteoarthr. Cartil. 21, 1933–1941 (2013).

8. A. P. Hollander, I. Pidoux, A. Reiner, C. Rorabeck, R. Bourne, A. R. Poole, Damage to type II collagen in aging and osteoarthritis starts at the articular surface, originates around chondrocytes, and extends into the cartilage with progressive degeneration. J. Clin. Invest. 96, 2859–2869 (1995).

9. V. D. Sree, A. B. Tepole, Computational systems mechanobiology of growth and remodeling: Integration of tissue mechanics and cell regulatory network dynamics. Curr. Opin. Biomed. Eng. 15 (2020), pp. 75–80.

10. J. Nam, B. D. Aguda, B. Rath, S. Agarwal, Biomechanical thresholds regulate inflammation through the NF-$\kappa$B pathway: Experiments and modeling. PLoS One. 4 (2009), doi:10.1371/journal.pone.0005262.

11. V. B. Shim, P. J. Hunter, P. Pivonka, J. W. Fernandez, A multiscale framework based on the Physiome markup languages for exploring the initiation of osteoarthritis at the bone-cartilage interface. IEEE Trans. Biomed. Eng. 58, 3532–3536 (2011).

12. G. A. Orozco, P. Tanska, C. Florea, A. J. Grodzinsky, R. K. Korhonen,

A novel mechanobiological model can predict how physiologically relevant dynamic loading causes proteoglycan loss in mechanically injured articular cartilage. Sci. Rep. 8 (2018), doi:10.1038/s41598-018-33759-3.

13. S. J. Millward-Sadler, D. M. Salter, Integrin-Dependent Signal Cascades in Chondrocyte Mechanotransduction. Ann. Biomed. Eng. 32, 435–446 (2004).

14. M. A. Wozniak, K. Modzelewska, L. Kwong, P. J. Keely, Focal adhesion regulation of cell behavior. Biochim. Biophys. Acta - Mol. Cell Res. 1692 (2004), pp. 103–119.

15. Z. Sun, S. S. Guo, R. Fässler, Integrin-mediated mechanotransduction. J. Cell Biol. 215 (2016), pp. 445–456.

16. R. F. Loeser, Integrins and chondrocyte-matrix interactions in articular cartilage. Matrix Biol. 39 (2014), pp. 11–16.

17. E. C. Arner, M. D. Tortorella, Signal transduction through chondrocyte integrin receptors induces matrix metalloproteinase synthesis and synergizes with interleukin-1. Arthritis Rheum. 38, 1304–1314 (1995).

18. N. Q. Balaban, U. S. Schwarz, D. Riveline, P. Goichberg, G. Tzur, I. Sabanay, D. Mahalu, S. Safran, A. Bershadsky, L. Addadi, B. Geiger, Force and focal adhesion assembly: A close relationship studied using elastic micropatterned substrates. Nat. Cell Biol. 3, 466–472 (2001).

19. J. Hoon, M. Tan, C.-G. Koh, The Regulation of Cellular Responses to Mechanical Cues by Rho GTPases. Cells. 5, 17 (2016).

20. K. H. Vining, D. J. Mooney, Mechanical forces direct stem cell behaviour in development and regeneration. Nat. Rev. Mol. Cell Biol. 18 (2017), pp. 728–742.

21. M. Amano, M. Nakayama, K. Kaibuchi, Rho-kinase/ROCK: A key regulator of the cytoskeleton and cell polarity. Cytoskeleton. 67 (2010), pp. 545–554.

22. G. Wang, A. Woods, S. Sabari, L. Pagnotta, L. A. Stanton, F. Beier, RhoA/ROCK Signaling Suppresses Hypertrophic Chondrocyte Differentiation. J. Biol. Chem. 279, 13205–13214 (2004).

23. M. Maekawa, T. Ishizaki, S. Boku, N. Watanabe, A. Fujita, A. Iwamatsu, T. Obinata, K. Ohashi, K. Mizuno, S. Narumiya, Signaling from Rho to the actin cytoskeleton through protein kinases ROCK and LIM-kinase. Science (80-. ). 285, 895–898 (1999).

24. D. R. Haudenschild, J. Chen, N. Pang, M. K. Lotz, D. D. D'Lima, Rho kinase-dependent activation of SOX9 in chondrocytes. Arthritis Rheum. 62, 191–200 (2010). 25. N. Wang, Review of cellular mechanotransduction. J. Phys. D. Appl. Phys. 50 (2017), p. 233002.

26. E. Lucchinetti, M. M. Bhargava, P. A. Torzilli, The effect of mechanical load on integrin subunits $\alpha 5$ and $\beta 1$ in chondrocytes from mature and immature cartilage

explants. Cell Tissue Res. 315, 385–391 (2004).

27. E. N. Olson, A. Nordheim, Linking actin dynamics and gene transcription to drive cellular motile functions. Nat. Rev. Mol. Cell Biol. 11 (2010), pp. 353–365.

28. C. S. Hill, J. Wynne, R. Treisman, The Rho family GTPases RhoA, Racl, and CDC42Hsregulate transcriptional activation by SRF. Cell. 81, 1159–1170 (1995).

29. S. Dupont, L. Morsut, M. Aragona, E. Enzo, S. Giulitti, M. Cordenonsi, F. Zanconato, J. Le Digabel, M. Forcato, S. Bicciato, N. Elvassore, S. Piccolo, Role of YAP/TAZ in mechanotransduction. Nature. 474, 179–184 (2011).

30. Y. Deng, A. Wu, P. Li, G. Li, L. Qin, H. Song, K. K. Mak, Yap1 Regulates Multiple Steps of Chondrocyte Differentiation during Skeletal Development and Bone Repair. Cell Rep. 14, 2224–2237 (2016).

31. Y. Deng, J. Lu, W. Li, A. Wu, X. Zhang, W. Tong, K. K. Ho, L. Qin, H. Song, K. K. Mak, Reciprocal inhibition of YAP/TAZ and NF-$\kappa$B regulates osteoarthritic cartilage degradation. Nat. Commun. 9, 1–14 (2018).

32. W. Zhong, Y. Li, L. Li, W. Zhang, S. Wang, X. Zheng, YAP-mediated regulation of the chondrogenic phenotype in response to matrix elasticity. J. Mol. Histol. 44, 587–595 (2013).

33. Y. Guan, X. Yang, W. Yang, C. Charbonneau, Q. Chen, Mechanical activation of mammalian target of rapamycin pathway is required for cartilage development. FASEB J. 28, 4470–4481 (2014).

34. B. Pal, H. Endisha, Y. Zhang, M. Kapoor, mTOR: A Potential Therapeutic Target in Osteoarthritis? Drugs R D. 15 (2015), pp. 27–36.

35. B. Yan, Z. Zhang, D. Jin, C. Cai, C. Jia, W. Liu, T. Wang, S. Li, H. Zhang, B. Huang, P. Lai, H. Wang, A. Liu, C. Zeng, D. Cai, Y. Jiang, X. Bai, mTORC1 regulates PTHrP to coordinate chondrocyte growth, proliferation and differentiation. Nat. Commun. 7 (2016), doi:10.1038/ncomms11151.

36. Q. Dai, Z. Xu, X. Ma, N. Niu, S. Zhou, F. Xie, L. Jiang, J. Wang, W. Zou, MTOR/Raptor signaling is critical for skeletogenesis in mice through the regulation of Runx2 expression. Cell Death Differ. 24, 1886–1899 (2017).

37. K. L. Posey, F. Coustry, A. C. Veerisetty, M. G. Hossain, M. J. Gambello, J. T. Hecht, Novel mTORC1 Mechanism Suggests Therapeutic Targets for COMPopathies. Am. J. Pathol. 189, 132–146 (2019).

38. L. Hui, V. Rodrik, R. M. Pielak, S. Knirr, Y. Zheng, D. A. Foster, mTOR-dependent suppression of protein phosphatase 2A is critical for phospholipase D survival signals in human breast cancer cells. J. Biol. Chem. 280, 35829–35835 (2005).

39. A. L. McNulty, H. A. Leddy, W. Liedtke, F. Guilak, TRPV4 as a therapeutic target for joint diseases. Naunyn. Schmiedebergs. Arch. Pharmacol. 388 (2015),

pp. 437–450.

40. C. J. O'Conor, H. A. Leddy, H. C. Benefield, W. B. Liedtke, F. Guilak, TRPV4-mediated mechanotransduction regulates the metabolic response of chondrocytes to dynamic loading. Proc. Natl. Acad. Sci. U. S. A. 111, 1316–1321 (2014).

41. M. N. Phan, H. A. Leddy, B. J. Votta, S. Kumar, D. S. Levy, D. B. Lipshutz, H. L. Suk, W. Liedtke, F. Guilak, Functional characterization of TRPV4 as an osmotically sensitive ion channel in porcine articular chondrocytes. Arthritis Rheum. 60, 3028–3037 (2009).

42. C. J. O'Conor, S. Ramalingam, N. A. Zelenski, H. C. Benefield, I. Rigo, D. Little, C. L. Wu, D. Chen, W. Liedtke, A. L. McNulty, F. Guilak, Cartilage-Specific Knockout of the Mechanosensory Ion Channel TRPV4 Decreases Age-Related Osteoarthritis. Sci. Rep. 6, 1–10 (2016).

43. S. Muramatsu, M. Wakabayashi, T. Ohno, K. Amano, R. Ooishi, T. Sugahara, S. Shiojiri, K. Tashiro, Y. Suzuki, R. Nishimura, S. Kuhara, S. Sugano, T. Yoneda, A. Matsuda, Functional gene screening system identified TRPV4 as a regulator of chondrogenic differentiation. J. Biol. Chem. 282, 32158–32167 (2007).

44. M. Lv, Y. Zhou, X. Chen, L. Han, L. Wang, X. L. Lu, Calcium signaling of in situ chondrocytes in articular cartilage under compressive loading: Roles of calcium sources and cell membrane ion channels. J. Orthop. Res. 36, 730–738 (2018).

45. E. Smedler, P. Uhlén, Frequency decoding of calcium oscillations. Biochim. Biophys. Acta - Gen. Subj. 1840, 964–969 (2014).

46. X. Gong, W. Xie, B. Wang, L. Gu, F. Wang, X. Ren, C. Chen, L. Yang, Altered spontaneous calcium signaling of in situ chondrocytes in human osteoarthritic cartilage. Sci. Rep. 7, 1–12 (2017).

47. N. Trompeter, J. Gardinier, V. DeBarros, M. Boggs, V. Gangadharan, W. Cain, L. Hurd, R. Duncan, bioRxiv, in press, doi:10.1101/2020.03.10.985713.

48. K. Novakofski, A. Boehm, L. Fortier, The small GTPase Rho mediates articular chondrocyte phenotype and morphology in response to interleukin-1$\alpha$ and insulin-like growth factor-1. J. Orthop. Res. 27, 58–64 (2009).

49. N. D. Leipzig, S. V. Eleswarapu, K. A. Athanasiou, The effects of TGF-$\beta$1 and IGF-I on the biomechanics and cytoskeleton of single chondrocytes. Osteoarthr. Cartil. 14, 1227–1236 (2006).

50. S. Pritchard, F. Guilak, Effects of interleukin-1 on calcium signaling and the increase of filamentous actin in isolated and in situ articular chondrocytes. Arthritis Rheum. 54, 2164–2174 (2006).

51. M. P. Vincenti, C. E. Brinckerhoff, Early response genes induced in chondrocytes stimulated with the inflammatory cytokine interleukin-1$\beta$. Arthritis Res. 3,

381–388 (2001).

52. Y. Y. Chow, K. Y. Chin, The Role of Inflammation in the Pathogenesis of Osteoarthritis. Mediators Inflamm. 2020 (2020), , doi:10.1155/2020/8293921.

53. M. B. Goldring, M. Otero, Inflammation in osteoarthritis. Curr. Opin. Rheumatol. 23 (2011), pp. 471–478.

54. M. Millerand, F. Berenbaum, C. Jacques, Danger signals and inflammaging in osteoarthritis. Clin. Exp. Rheumatol. 37 (2019), pp. 48–56.

55. T. Sillat, G. Barreto, P. Clarijs, A. Soininen, M. Ainola, J. Pajarinen, M. Korhonen, Y. T. Konttinen, R. Sakalyte, M. Hukkanen, P. Ylinen, D. C. E. Nordström, Toll-like receptors in human chondrocytes and osteoarthritic cartilage. Acta Orthop. 84, 585–592 (2013).

56. A. Weber, P. Wasiliew, M. Kracht, Interleukin-1 (IL-1B) pathway. Sci. Signal. 3 (2010), p. 1.

57. C. Lianxu, J. Hongti, Y. Changlong, NF-$\kappa$Bp65-specific siRNA inhibits expression of genes of COX-2, NOS-2 and MMP-9 in rat IL-1$\beta$-induced and TNF-$\alpha$-induced chondrocytes. Osteoarthr. Cartil. 14, 367–376 (2006).

58. S. Madhavan, M. Anghelina, D. Sjostrom, A. Dossumbekova, D. C. Guttridge, S. Agarwal, Biomechanical Signals Suppress TAK1 Activation to Inhibit NF-$\kappa$B Transcriptional Activation in Fibrochondrocytes. J. Immunol. 179, 6246–6254 (2007).

59. Choi, Jo, Park, Kang, Park, NF-B Signaling Pathways in Osteoarthritic Cartilage Destruction. Cells. 8, 734 (2019).

60. R. M. Clancy, P. F. Gomez, S. B. Abramson, Nitric oxide sustains nuclear factor kappaB activation in cytokine-stimulated chondrocytes. Osteoarthr. Cartil. 12, 552–558 (2004).

61. J. N. Sharma, A. Al-Omran, S. S. Parvathy, Role of nitric oxide in inflammatory diseases. Inflammopharmacology. 15 (2007), pp. 252–259.

62. K. Vuolteenaho, T. Moilanen, U. Jalonen, A. Lahti, R. Nieminen, H. M. Van Beuningen, P. M. Van Der Kraan, E. Moilanen, TGF$\beta$ inhibits IL-1 -induced iNOS expression and NO production in immortalized chondrocytes. Inflamm. Res. 54, 420–427 (2005).

63. B. Ma, L. Zhong, C. A. Van Blitterswijk, J. N. Post, M. Karperien, T cell factor 4 is a pro-catabolic and apoptotic factor in human articular chondrocytes by potentiating nuclear factor $\kappa$B signaling. J. Biol. Chem. 288, 17552–17558 (2013).

64. C. Malemud, Negative Regulators of JAK/STAT Signaling in Rheumatoid Arthritis and Osteoarthritis. Int. J. Mol. Sci. 18, 484 (2017).

65. R. C. Olney, K. Tsuchiya, D. M. Wilson, M. Mohtai, W. J. Maloney, D. J. Schurman, R. L. Smith, Chondrocytes from osteoarthritic cartilage have increased

expression of insulin-like growth factor I (IGF-I) and IGF-binding protein-3 (IGFBP-3) and -5, but not IGF-II or IGFBP-4. J. Clin. Endocrinol. Metab. 81, 1096–1103 (1996).

66. J. Martel-Pelletier, J. A. Di Battista, D. Lajeunesse, J. P. Pelletier, IGF/IGFBP axis in cartilage and bone in osteoarthritis pathogenesis. Inflamm. Res. 47 (1998), pp. 90–100.

67. R. C. Olney, D. M. Wilson, M. Mohtai, P. J. Fielder, R. L. Smith, Interleukin-1 and tumor necrosis factor-$\alpha$ increase insulin-like growth factor-binding protein-3 (IGFBP-3) production and IGFBP-3 protease activity in human articular chondrocytes. J. Endocrinol. 146, 279–286 (1995).

68. A. M. Grabiec, C. Angiolilli, L. M. Hartkamp, L. G. M. Van Baarsen, P. P. Tak, K. A. Reedquist, JNK-dependent downregulation of FoxO1 is required to promote the survival of fibroblast-like synoviocytes in rheumatoid arthritis. Ann. Rheum. Dis. 74, 1763–1771 (2015).

69. T. Matsuzaki, O. Alvarez-Garcia, S. Mokuda, K. Nagira, M. Olmer, R. Gamini, K. Miyata, Y. Akasaki, A. I. Su, H. Asahara, M. K. Lotz, FoxO transcription factors modulate autophagy and proteoglycan 4 in cartilage homeostasis and osteoarthritis. Sci. Transl. Med. 10, eaan0746 (2018).

70. J. Deschner, B. Rath-Deschner, S. Agarwal, Regulation of matrix metalloproteinase expression by dynamic tensile strain in rat fibrochondrocytes. Osteoarthr. Cartil. 14, 264–272 (2006).

71. A. Dossumbekova, M. Anghelina, S. Madhavan, L. He, N. Quan, T. Knobloch, S. Agarwal, Biomechanical signals inhibit IKK activity to attenuate NF-$\kappa$B transcription activity in inflamed chondrocytes. Arthritis Rheum. 56, 3284–3296 (2007).

72. B. Fermor, J. Brice Weinberg, D. S. Pisetsky, M. A. Misukonis, A. J. Banes, F. Guilak, The effects of static and intermittent compression on nitric oxide production in articular cartilage explants. J. Orthop. Res. 19, 729–737 (2001).

73. B. Fermor, J. B. Weinberg, D. S. Pisetsky, F. Guilak, The influence of oxygen tension on the induction of nitric oxide and prostaglandin E2 by mechanical stress in articular cartilage. Osteoarthr. Cartil. 13, 935–941 (2005).

74. J. Martel-Pelletier, A. J. Barr, F. M. Cicuttini, P. G. Conaghan, C. Cooper, M. B. Goldring, S. R. Goldring, G. Jones, A. J. Teichtahl, J. P. Pelletier, Osteoarthritis. Nat. Rev. Dis. Prim. 2 (2016), pp. 1–18.

75. L. G. Alexopoulos, L. A. Setton, F. Guilak, The biomechanical role of the chondrocyte pericellular matrix in articular cartilage. Acta Biomater. 1, 317–325 (2005). 76. A. Erdemir, Open knee: A pathway to community driven modeling and simulation in joint biomechanics. J. Med. Devices, Trans. ASME. 7, 1 (2013).

77. S. Mukherjee, R. Lesage, W. Wilson, L. Geris. Multiscale modeling to inves-

tigate the role of mechanical loading in articular cartilage. 14th World Congress in Computational Mechanics (WCCM) and ECCOMAS Congress 2020, 11-15 January 2021, Virtual Congress. (2021).

78. P. Julkunen, W. Wilson, J. S. Jurvelin, R. K. Korhonen, Composition of the pericellular matrix modulates the deformation behaviour of chondrocytes in articular cartilage under static loading. Med. Biol. Eng. Comput. 47, 1281–1290 (2009).

79. A. A. Young, S. McLennan, M. M. Smith, S. M. Smith, M. A. Cake, R. A. Read, J. Melrose, D. H. Sonnabend, C. R. Flannery, C. B. Little., Proteoglycan 4 downregulation in a sheep meniscectomy model of early osteoarthritis. Arthritis Res. Ther. 8 (2006), doi:10.1186/ar1898.

80. J. B. Thorlund, A. Holsgaard-Larsen, M. W. Creaby, G. M. Jørgensen, N. Nissen, M. Englund, L. S. Lohmander, Changes in knee joint load indices from before to 12 months after arthroscopic partial meniscectomy: A prospective cohort study. Osteoarthr. Cartil. 24, 1153–1159 (2016).

81. S. Fu, H. Meng, S. Inamdar, B. Das, H. Gupta, W. Wang, C. L. Thompson, M. M. Knight, Activation of TRPV4 by mechanical, osmotic or pharmaceutical stimulation is anti-inflammatory blocking IL-1$\beta$ mediated articular cartilage matrix destruction. Osteoarthr. Cartil. 29, 89–99 (2021).

82. R. Potla, M. Hirano-Kobayashi, H. Wu, H. Chen, A. Mammoto, B. D. Matthews, D. E. Ingber, Molecular mapping of transmembrane mechanotransduction through the $\beta$1 integrin–CD98hc–TRPV4 axis. J. Cell Sci. 133 (2020), doi:10.1242/jcs.248823.

83. C. Ji, C. A. McCulloch, TRPV4 integrates matrix mechanosensing with Ca2+ signaling to regulate extracellular matrix remodeling. FEBS J. (2020), , doi:10.1111/febs.15665.
